# MAGE: Monte Carlo method for Aberrant Gene Expression

**DOI:** 10.1101/2024.11.08.622686

**Authors:** Matthew Beltran, Richard I. Joh

**Affiliations:** Department of Physics, Virginia Commonwealth University, Richmond, VA 23220; Massey Cancer Center, Virginia Commonwealth University, Richmond, VA 20220

## Abstract

Identifying genes that are aberrantly expressed is an important first step in the diagnosis and treatment of many diseases. Conventionally, differential expression (DE) analysis is used to screen gene expression profiles to identify functionally associated genes. DE often relies on the variance and fold change in expression from individual genes, which does not consider the expression of all other genes within the profile. When the overall gene expression is skewed, DE does not capture outliers in gene expression. To address this, we have developed a non-parametric DE method based on the probability density for an entire expression profile to select genes that deviate from the global distribution between two gene expression profiles with multiple replicates. Rather than assuming a particular distribution of expression per gene, our method assumes that aberrantly expressed genes (AEGs) will exhibit expression patterns distinguishable from non-AGEs which make up the majority of the profile.

Here we introduce our nonparametric method (MAGE: Monte Carlo method for aberrant gene expression) and demonstrate that MAGE can identify AEGs that are not found by conventional DE analyses. The main feature of MAGE is (1) identifying outliers based on the expression profile of all genes rather than performing DE analyses on a per-gene basis and (2) consideration of the variance in expression between two different conditions. We also compared our results with traditional DE analysis as well as density-based clustering methods. MAGE produces consistent results in a variety of conditions and performs conservatively with the addition of noise. We also applied MAGE to single-cell RNA-seq samples and demonstrated that the analysis is robust with subsampling.

## Introduction

One of the first steps for tackling human diseases is to understand the molecular players behind the emergence of diseases, and disease-associated genes can be further studied as potential therapeutic targets. A common practice is to utilize expression profile data (often RNA-seq) to identify differentially expressed genes (DEG) by comparing differences in expression levels between healthy and diseased tissues or control and treatment samples [1–3]. When a gene’s expression is significantly different from that of the control, it is considered to be differentially expressed, and DEGs can become candidate biomarkers for diagnosis and prognosis of human diseases. Additionally, the transcriptional, translational, and post-translational regulation of DEGs can be further targeted for potential use in drug development.

Most methods for DEG identification fall in the category of parametric statistical significance tests, where gene expression levels are assumed to be from a known probability distribution described by several parameters [2,4,5]. Since expression levels are typically measured by read counts from RNA-seqs, discrete probability distributions (e.g. Poisson [6,7] and negative-binomial [4,5] for DESeq and EdgeR) are commonly employed to characterize expression profiles. The Poisson distribution, as governed by a single parameter, is preferred for its simplicity, however, it is limited by the constraint that the variance and mean are equal. This is often not applicable as the biological variation among distinct replicates is larger than the expected technical variation of sample preparation [8]. Because of this constraint, the Poisson distribution often does not fully account for the deviation in many expression profiles and may result in higher false-positive discovery rates than other probability distributions. While the negative-binomial distribution (ND) includes separate parameters for the mean and variance, the number of samples is sometimes too small to effectively evaluate both parameters. With each DE method based on a different set of assumptions, this has led to a large number of context-dependent health- and disease-associated DEGs [9], and challenges exist in devising methods for narrowing the large number of DEGs robustly. Prior studies suggest different genes may not conform to the same distribution, and an approach that can handle many types of probability distributions and extract biologically relevant gene signatures is needed [10].

Our approach takes inspiration from several classes of nonparametric machine learning-based methods previously used to identify DEGs. Namely, Gaussian mixture models (GMM) which have been used for outlier detection in RNA-seq datasets [11,12], operate on a similar premise of considering the underlying probability density function (PDF) from each gene. Other methods of outlier detection involve distance-based clustering methods such as k-nearest neighbor [13], and density-based approaches such as DBSCAN [14,15]. Information entropy is also used to discover cluster genes in a noise-resistant manner [16]. Previous algorithms have been developed for identifying outlier genes among RNA-seq profiles. One example to identify aberrant gene expression in rare disorders is OUTRIDER [17], which utilizes an auto-encoder to account for unknown covariation between genes, followed by a statistical p-value determined using ND. Similarly, ABEILLE [18] utilizes a variational auto-encoder and introduces an anomaly score that is determined using an isolation forest approach. This removes the distributional assumptions used in OUTRIDER and other parametric DEG methods, however, it introduces the need to use multiple predetermined thresholds for AE classification. ORdensity is another algorithm used to find outlier genes reliably from microarray data [19].

With this in mind, we have developed a method that identifies AEGs by estimating the 2-dimensional (2D) PDF of each gene and comparing it with the combined 2D cumulative PDF (CPDF) from all genes. We assume that true AEGs will exhibit deviated expression compared to those of all others. It is also vital to consider the computational costs of conducting complex analysis for the identification of DEGs due to the sheer size of the omics datasets that are frequently encountered. Our method performs robustly while remaining computationally cost-effective so that it can be applied to expression profiles containing thousands of genes across many samples.

## Methods

### Aberrant expression vs differential expression

It is important to consider the similarities and distinctions between traditional DEGs and AEGs. Both are interested in the outlier genes within an expression profile, however, DEG is assessed on an individual gene basis, while AEG is based on the overall gene expression patterns across two different conditions. In many cases, there is a significant overlap between DEGs and AEGs, and we show that this overlap is especially evident in simple expression profiles examining small effects between conditions. We have observed that as the expression behavior becomes more complex, the agreement between DEGs and AEGs decreases.

### Setting up MAGE

The input data are expression profiles from two different sample conditions (e.g. cell/tissue types, diagnosis, treatment/control pairs, etc.) each of which contains *n* number of biological replicates, and *n* must be greater than 2. Figure 1 illustrates the overview of MAGE and its workflow. MAGE does not have built-in normalization, so the input expression profile should be in pre-normalized units such as TPM, RPKM, etc. to account for differences in transcript length and sequencing depth. Further filtering can be performed by removing genes with near-zero reads in a majority of samples. Filtering of low read count genes is recommended to significantly reduce processing times and avoid noisy expression. Similar to other methods of finding gene signatures, our algorithm can produce results for any profiles with two conditions and more than one replicate sample per condition.

**Fig 1.**
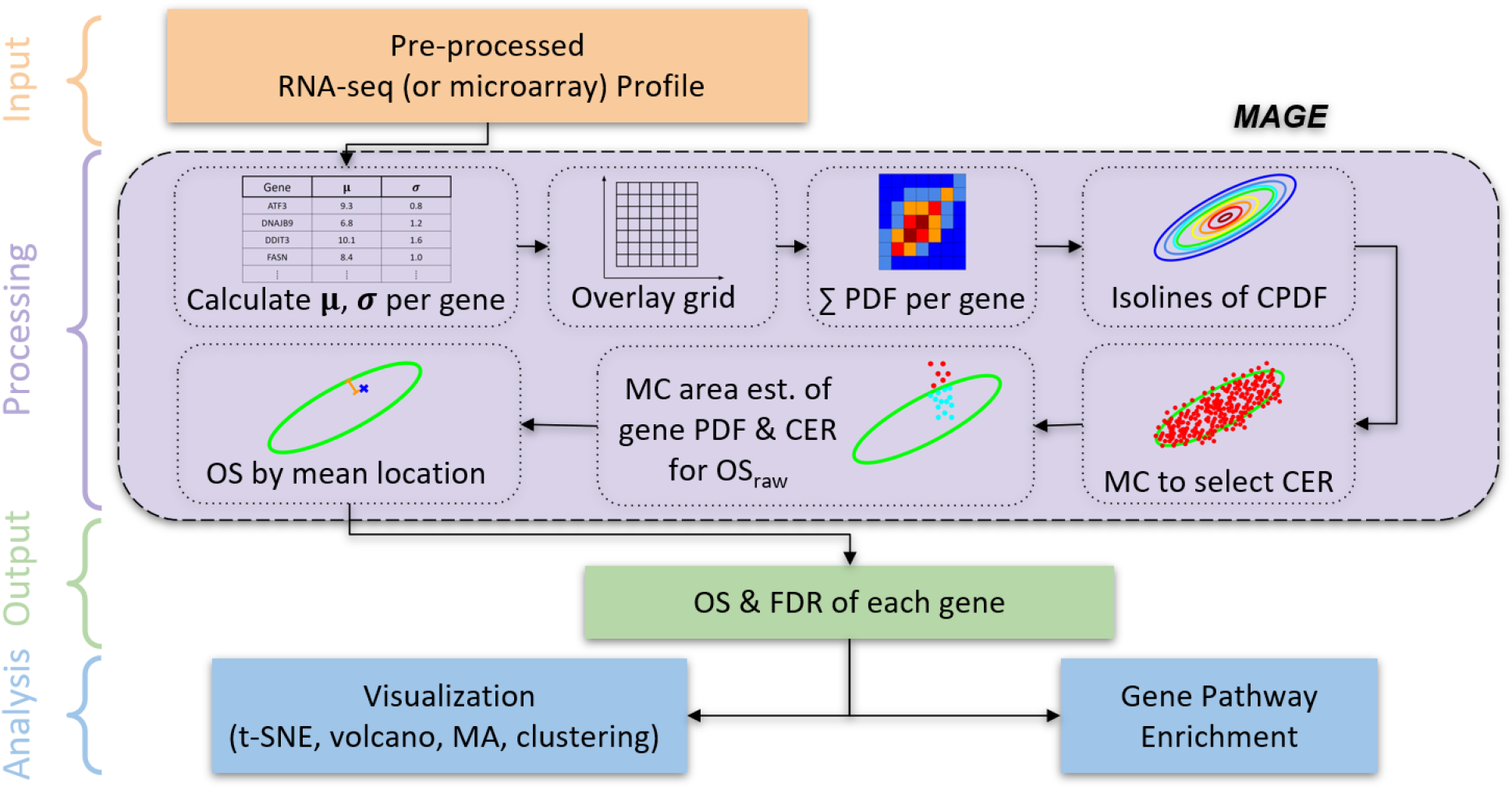
MAGE analysis overview. MAGE accepts RNA-seq or microarray profiles from two conditions and at least two replicates per condition. MAGE provides output as outlier scores based on the cumulative PDF of all genes and the distance between the CER and the mean coordinate of each gene. MC sampling is performed twice to determine CER from gene expression profiles and to quantify the outlier score and FDR for individual genes. Further analysis includes visualization techniques and gene pathway enrichment to identify the functional relevance of AEGs.

First, we calculate the mean and standard deviation (SD) from the expression of a gene at a given condition (eqs. 1-2). Then we calculate the probability density function (PDF) over the 2D expression region (eqs. 3-5) and estimate the cumulative sum of PDFs of individual genes (eqs. 6-7). The resulting cumulative probability density function (CPDF) approximates a two-dimensional PDF or density matrix for the whole profile, and CPDF is evaluated at fixed indices within a grid that overlays the expression. We choose the range of the grid to cover beyond the mean expression values by adding four times the average SD of each gene to the respective grid range. The height and width of each grid are determined by dividing the adjusted ranges by the number of indices along one dimension (eq. 5). The number of grid indices is an adjustable parameter that can be increased for higher precision or decreased for faster computation. Figure S1 shows CPDFs from selected individual genes while varying the total number of genes, and Figure S2 displays the entire CPDF (*N* ≅ 15,000). It is possible that given a large enough number of indices, each index may have very few contributing genes. To avoid this, the total number of indices should be significantly less than the number of genes, and the range of the grid should be set high enough to fully capture the data structure.

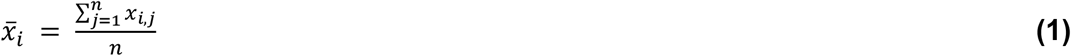

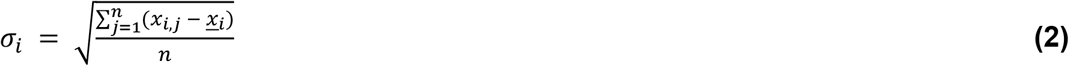

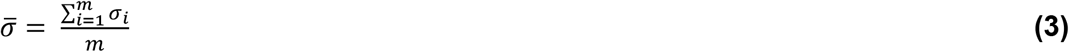

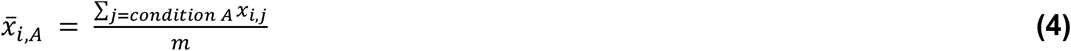

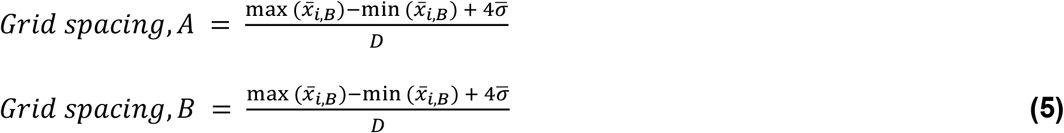

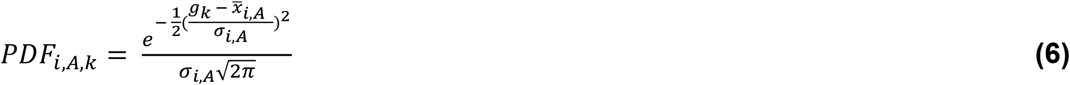

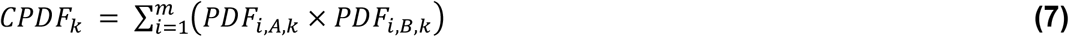

*x*_*i,j*_ is the read count of the i^th^ gene in the j^th^ sample, and 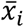 represents the average expression of the i^th^ gene. *n* and *m* represent the number of replicate samples and the number of genes, respectively. *σ*_*i*_ is the standard deviation of i^th^ gene across replicates, and 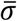 is the mean standard deviation of all genes. 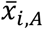 is the mean expression of the i^th^ gene in condition A. Eq. 5 denotes the selection of grid space based on the maximum and minimum expression in each condition A or B. *D*^2^ is the total number of grid indices, and *g*_*k*_ is the location of grid index k. *PDF*_*i,A,k*_ is the Gaussian probability density function of the ith gene in condition A at grid index k, while *CPDF*_*k*_ is the cumulative probability density of all genes evaluated at grid index k.

### Determining the characteristic expression region (CER)

From the probability density matrix, we generate a contour plot to find isolines across the CPDF. We define CER as an optimized contour of CPDF, which captures the gene expression profile of the majority of genes. CER from the CPDFs can be determined using contour lines (Figure S3), and we implemented an iterative optimization algorithm to find the optimal contour level as the CER. First, we select a fixed number of contour lines, which is an adjustable parameter. For each contour line, we quantify contour effectiveness (*CE*) by the fraction of enclosed genes, and the contour line that matches the target containment fraction is CER. CER can consist of multiple unconnected components (Figure S4). To calculate the actual fraction of CPDF containment, we perform Monte Carlo (MC) sampling (10 points per gene) from each gene’s PDF (Figure 2A, S3B). Since most profiles contain thousands of genes, sampling only a few points from each gene’s PDF still results in a significant (> 10,000) sample size for area estimation. We quantify the total fraction of MC points inside a given contour. Then we compare the actual containment fraction (A) with the target containment fraction (T) (eq. 8). The target containment fraction is a user-set parameter, and we use 95%. *CE* is quantified as

**Fig 2.**
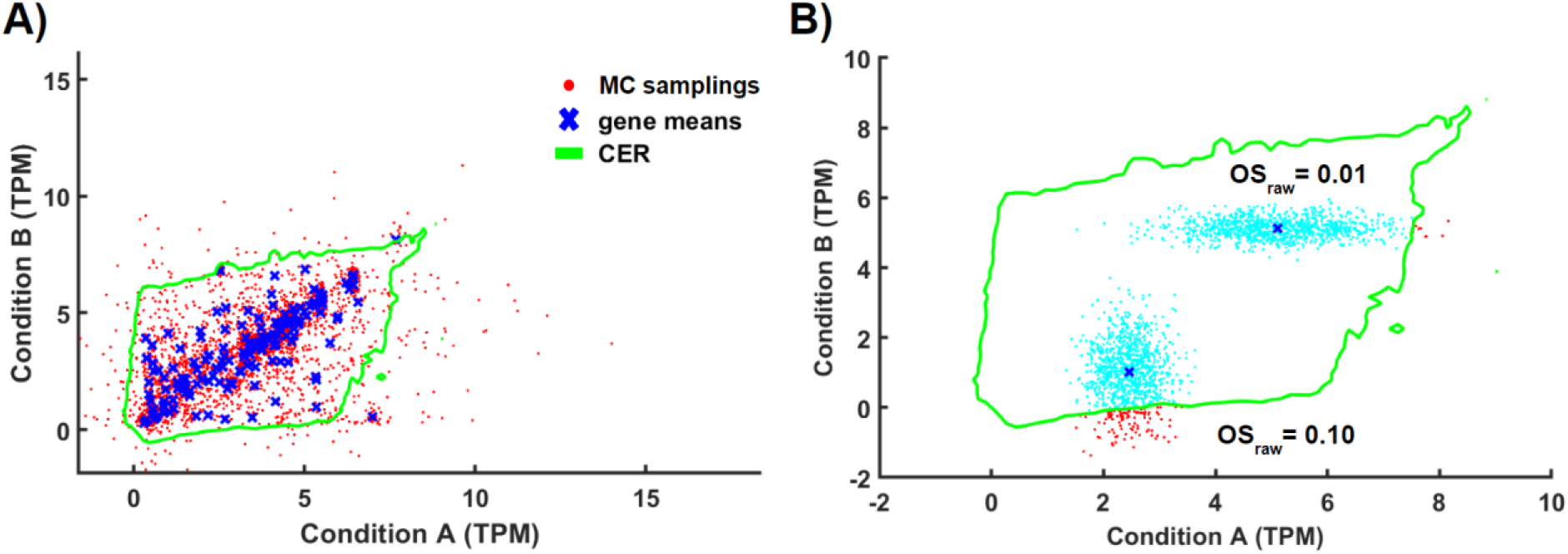
Selection of the characteristic expression region (CER) and estimation of *OS*_*raw*_. (A) For a given contour, Monte Carlo (MC) sampling (fixed number *N*_*mc*1_) is performed for each gene from its 2D PDF. Then we calculate the fraction of all MC sampled points (total *N*_*gene*_ × *N*_*mc*1_ points) inside the contour. Blue and red points represent gene means and MC sampled points, respectively. CER is selected to match the target fraction of MC sampled points lying inside. (B) For individual genes, further points (*N*_*mc*2_ per gene) are sampled from the individual gene PDF. The fraction of these points outside CER (denoted red) determines *OS*_*raw*_.

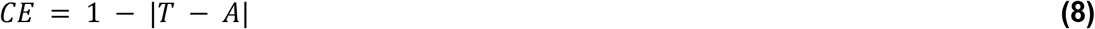

where *T*, and *A* are the target fraction of containment, the actual fraction of containment, and the size (area) of the contour relative to the size of the grid, respectively. Figure S5 illustrates the performance of CER and other isolines.

### Quantification of raw outlier score (*OS*_*raw*_)

We consider genes that are likely to be found outside the CER to be aberrantly expressed, which we term AEGs. To quantify this, we implemented a second round of MC sampling. The expression of each gene follows its PDF, and CER can have an arbitrary shape. A larger number of points (1000 in this study as an adjustable parameter) are sampled from each gene’s PDF. The ratio of points falling inside and outside of the CER is quantified as the raw outlier score (*OS*_*raw*_) (Figure 2B). For an individual gene, *OS*_*raw*_ represents the probability of that gene’s expression to be outside of the CER as

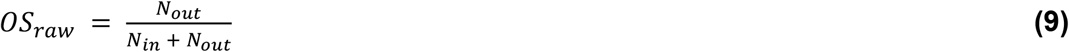

where *P* is the number of sampled points inside/outside of CER.

### Adjusting high variance genes based on mean location

Two genes may have the same outlier probability, but the variance of one gene is greater than that of the other. Simple quantification of *OS*_*raw*_ does not account for the degree of certainty affected by the variance of points. Figure S6 illustrates two genes with similar *OS*_*raw*_ and significantly different variances in expression. A gene with very high variance can lead to significant *OS*_*raw*_ regardless of the sample mean values. To adjust for these differences in variance, we chose to adjust *OS*_*raw*_ since it is less certain that genes with high variance are meaningfully deviating from the CER. Since *OS*_*raw*_ has a maximum value of 1, we devised a ranked adjustment which also ranges from 0 to 1. We implement this by subtracting the adjustment from the *OS*_*raw*_ to determine *OS* (eq. 10). The correction is determined as

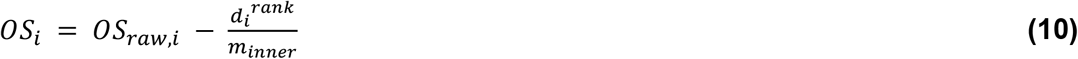

where *d*_*i*_^*rank*^ is the nearest distance from the gene mean expression to CER, and *m*_*inner*_ is the number of genes closer to the CER than the i^th^ gene. The adjustment is determined by finding the nearest distance from each gene (mean in both conditions) to the CER and rank-ordering all genes with a mean expression falling within the CER. Therefore, genes that are near the center of the CER will be penalized the most whereas genes near CER undergo a small reduction in *OS*. Figure S7 illustrates the change in *OS*, and the adjustment largely eliminates the effect of high variance on *OS*. Figure S8 shows that *OS* is adjusted due to the distance to CER, and genes interior to CER are unlikely to be called outliers after this adjustment.

### Filtering low/high expression *OS* values

Often there is a significant correlation of expression between 2 samples. Because our method considers the density of the expression probability, genes that are very lowly or highly expressed in both samples will receive a high *OS*_*raw*_. Housekeeping genes may fall into this category since their expression tends to be consistent across many conditions [20]. These genes may or may not be biologically relevant in the mechanism of interest. Therefore, we provide the option to ignore these genes by setting the *OS* for lowly/highly expressed genes to zero. The threshold is determined based on the target containment. For example, if the target containment is 95%, then the lowest 2.5%, and highest 2.5% genes will receive an *OS* of zero.

### FDR estimation

To estimate the rate of false discoveries in datasets where the true FDR is unknown, we used a standard permutation-based method by running MAGE while using permutated sample classes. Randomly assigning treatment/control labels guarantees a true null hypothesis and therefore any predicted gene signatures are considered falsely discovered. The ratio of false gene signatures to total gene signatures is found to determine the FDR (eq. 11-12). To avoid overestimation of FDR, we used the modification proposed by Xie et. al. of only using non-signature genes for the estimation of FDR [21,22].

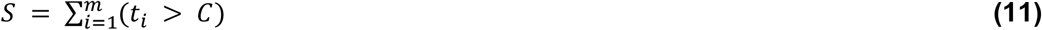

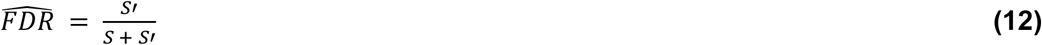

where *S* is the number of gene signatures. *t*_*i*_ and *C* the test statistic (FC, p-value, *OS*, etc.) for the i^th^ gene and the test statistic classification threshold, respectively. FDR estimation as a function of *OS* is shown in Figure S9.

### Effects of noise

RNA-seq data is susceptible to many sources of noise which can arise from errors during transcription or splicing, PCR amplification biases, and barcode swapping during library preparation [23–25]. Noise can either introduce false positives by elevating the test statistics across a broad set of genes, or true-positive genes may be missed as the statistical power is reduced. We tested the performance of MAGE against the standard 2 sample t-test to analyze the *γ* - T3 breast cancer profile with different amounts of Gaussian noise introduced (Figure S10). The noise was introduced by adding a normally distributed random number with SD (*σ*_n_) for each gene (eq. 13) as

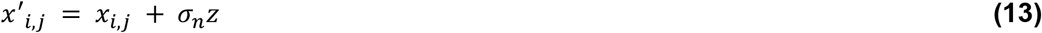

where *z* is a normally distributed random variable.

The effect of noise is shown by a widening of the CER (Figure S10), which results in an overall reduction in *OS*. Figures S11-S12 show the performance of MAGE and identified AEGs in two different datasets. DE analysis (as described below) identifies more DEGs with the presence of noise, but AEG identification remains conservative for small to moderate levels of noise introduced. These results suggest that MAGE performs well with noisy data.

### Density-based clustering vs MAGE

MAGE works similarly in principle to density-based classification methods, and we performed a side-by-side comparison between a widely used density-clustering algorithm, DBSCAN [14]. DBSCAN performs solely on individual data points without consideration for variance, and this constitutes a major downside in the analysis of gene expression as there is no way to incorporate multiple replicates to improve predictions. For the input to DBSCAN, we used the mean expression of each gene within both conditions. Figure 3 shows the comparison between MAGE and DBSCAN. DBSCAN can find the outlier without the consideration of sample variance whereas MAGE can identify interior genes within CER as AEGs depending on variance.

**Figure 3.**
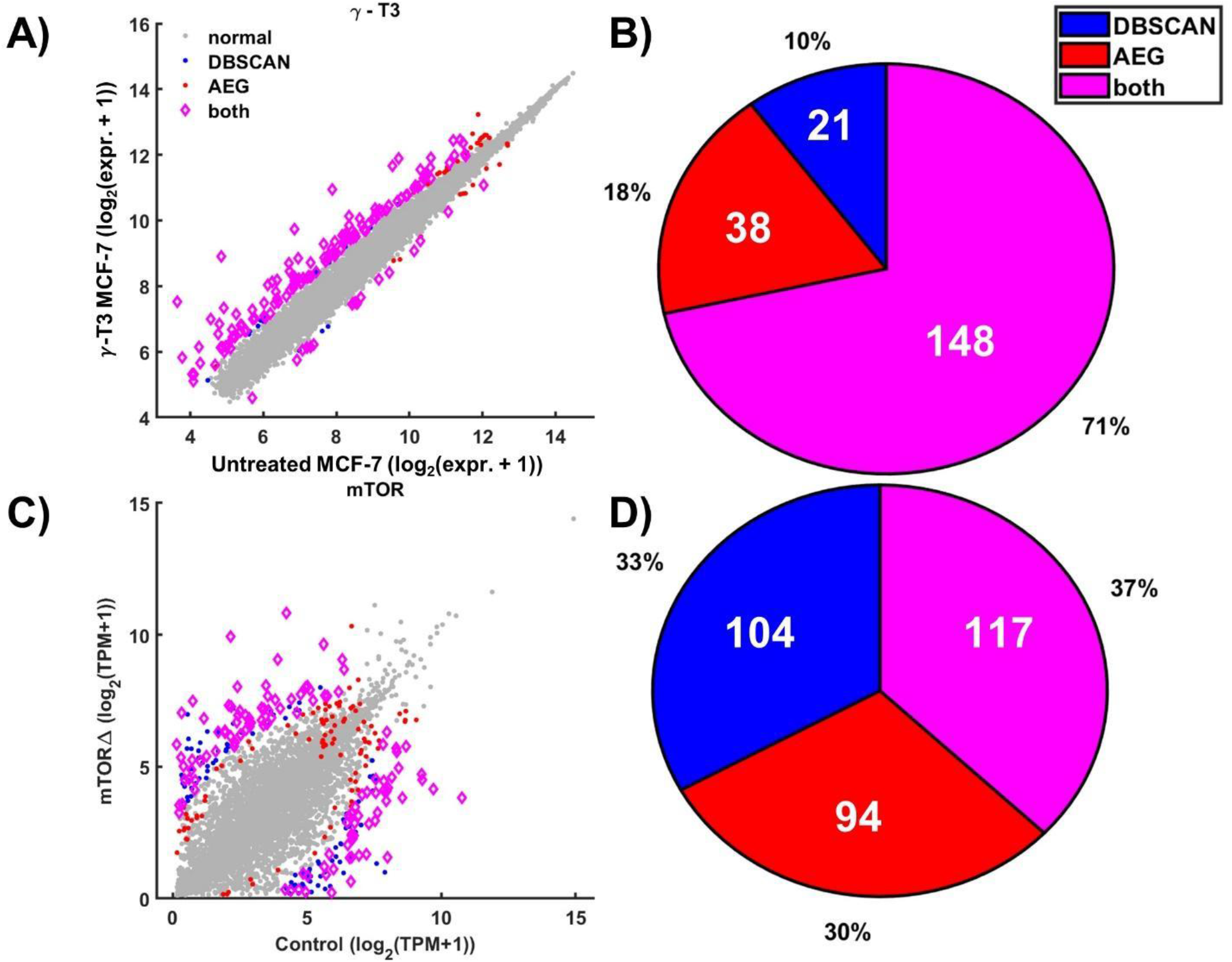
Comparison of DBSCAN and MAGE. (A) Gene mean values from the *γ*-T3 breast cancer profiles (data from GSE21946). Blue, red, and magenta marked points indicate genes identified by DBSCAN (*ε* = 0.3, *minpts* = 200), AEG (*OS* > 0.65), and both, respectively. (B) Overlap between DBSCAN and MAGE for *γ*-T3 breast cancer profile genes. (C) Gene mean values from the mTOR KO mouse profile (data are from GSE134316). Signature genes were identified by DBSCAN (*ε* = 0.7, *minpts* = 80), AEG (*OS* > 0.1). (D) Overlap between DBSCAN and MAGE for mTOR KO mouse profile genes.

### Pathway/GO enrichment

To evaluate the biological significance of identified signature genes, we performed gene set enrichment of the gene ontology (GO) biological process (BP) terms [26] and KEGG pathways [27] using DAVID [28]. Significantly enriched pathways were selected and sorted using the Benjamini and Hochberg false discovery rate to account for multiple tests [21].

### Identification of DEGs

To compare the performance of MAGE, we also identified DEGs by assessing the logarithmic fold-change (FC) along with a standard 2-sample t-test. The test statistic t was determined for each gene (eq. 14) and used to find the 2-tailed p-value representing the probability of a deviation in mean expression for a single gene across the 2 sample conditions. DEGs were selected by finding genes with a p-value below 0.05 and an FC above a specified threshold. P-values were also assessed and compared using edgeR [4].

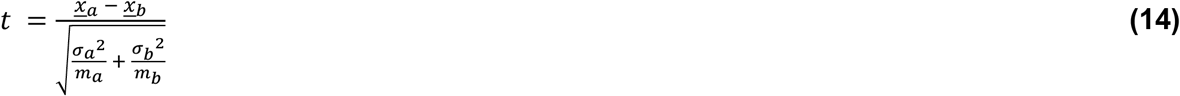

### Data collection and preparation

We used NCBI GEO expression profiles (GSE21946, GSE134316, GSE98816, and GSE99235). GSE21946 is a microarray dataset consisting of expression levels from 22,277 genomic loci in human breast cancer samples (MCF-7 cells) subjected to treatment with gamma-tocotrienol (*γ* - T3) [29]. We averaged the expression of multiple probes from the same gene, reducing the profile to 14,054 genes. Genes were filtered to ensure they contained non-zero expression in 6 out of 8 samples, leaving 13,639 genes for DE and AE analysis. *OS* and FDR values were determined using MAGE, and Figure 4 illustrates the CER and distribution of *OS*. Figure S9A shows the FDR as a function of *OS* threshold for AEG classification. GSE134316 contains 3 healthy control samples and 3 mTOR knockout samples from mouse bone marrow [30]. The initial set of 49,431 genes was filtered to remove genes containing zero expression in all samples, and the remaining 34,358 genes were log-shifted. GSE98816 and GSE99235 contain single-cell RNA-seq (scRNA-Seq) profiles of mouse blood vessel cells from the brain and lung respectively. We focused on only the two largest cell types common to both profiles, endothelial and mural cells. Both brain and lung profiles consist of roughly 20,000 genes with reads measured in 3,186 brain and 1,504 lung cells and were filtered to ensure they contained non-zero expression in at least 100 samples, leaving 10,461 genes for downstream analysis.

**Figure 4.**
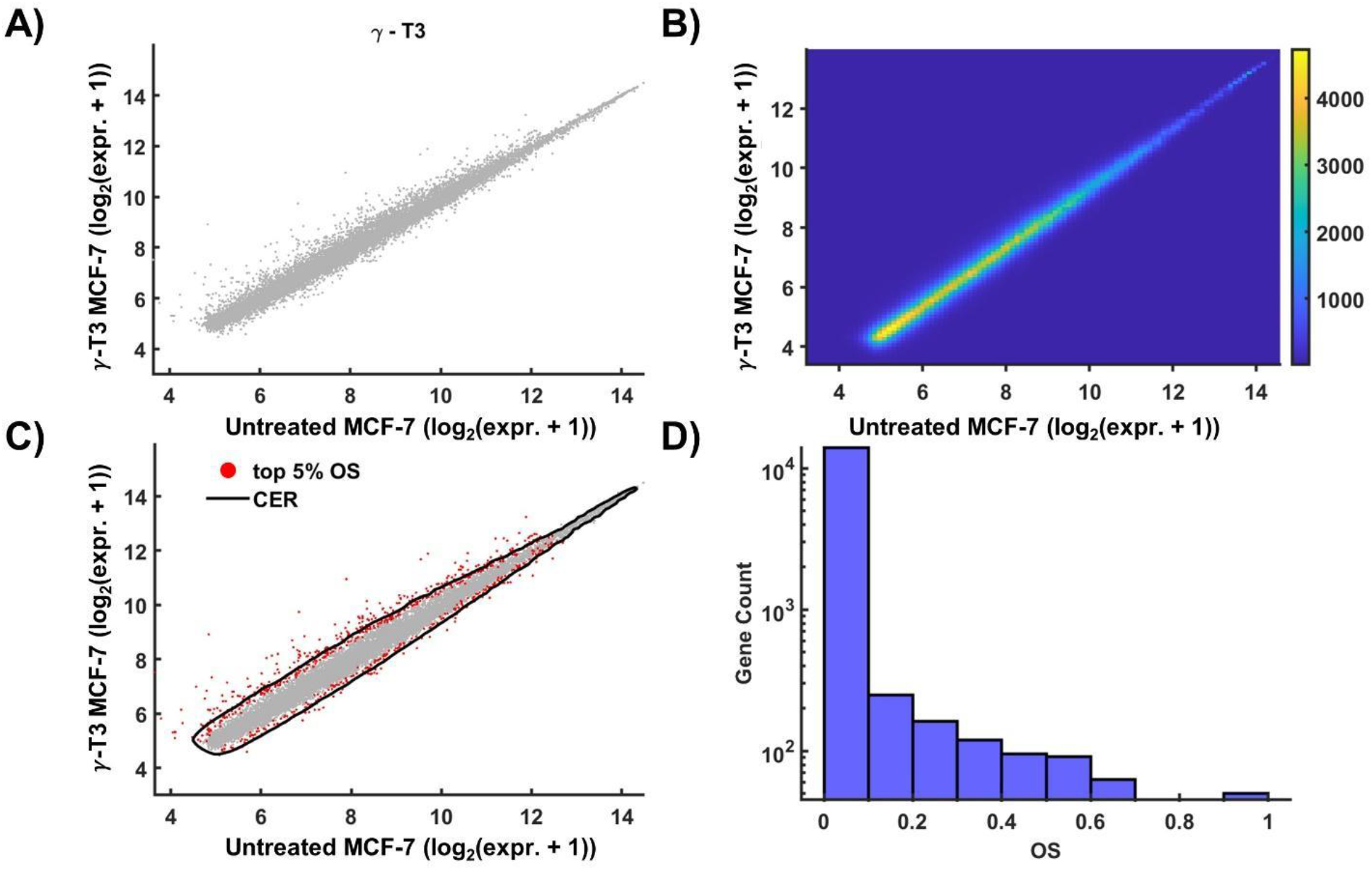
MAGE applied to the breast cancer *γ* – T3 treatment profile. (A) Mean RNA-seq TPM levels of each gene per sample type after filtering. (B) CPDF from all genes. (C) CER based on the CPDF. (D) The distribution of *OS* from 1,000 MC sampling per gene.

## Code availability

All data preparation, processing, and figures were performed in *MATLAB version R2021a* [31]. All code used in this study is available on GitHub github.com/beltranmm/MAGE

## Results

### Analysis of human breast cancer microarray data

To compare the performance of MAGE and the conventional DEG identification, we used both approaches to analyze a simple two-condition microarray expression profile. This dataset, obtained from the NCBI GEO (GSE21946), consists of human breast cancer samples (MCF-7 cells) subjected to treatment with gamma-tocotrienol (*γ* - T3) [29], an antioxidant and form of vitamin E known for its demonstrated inhibition of tumor growth in various types of cancers [32]. Figure 4 shows the identification of AEGs, which are distributed mostly outside of the CER, and the distribution of *OS*.

Figure 5 compares AEGs and DEGs based on FC, p-value, mean expression, and *OS*. We see significant AEG/DEG agreement since there is a correlation between FC and *OS* (Figure 5C). MAGE assigns higher *OS* to genes with higher mean expression (Figure 5B) while the conventional method favors genes with lower expression. The preference for selecting higher expressed genes is explained by the disproportionate number of lowly and highly expressed genes. Since our method considers CPDF of all genes, the higher expression region will naturally have a lower density, resulting in a narrowing of the CER (Figure 4C), and is, therefore, more likely to have a higher *OS*.

**Figure 5.**
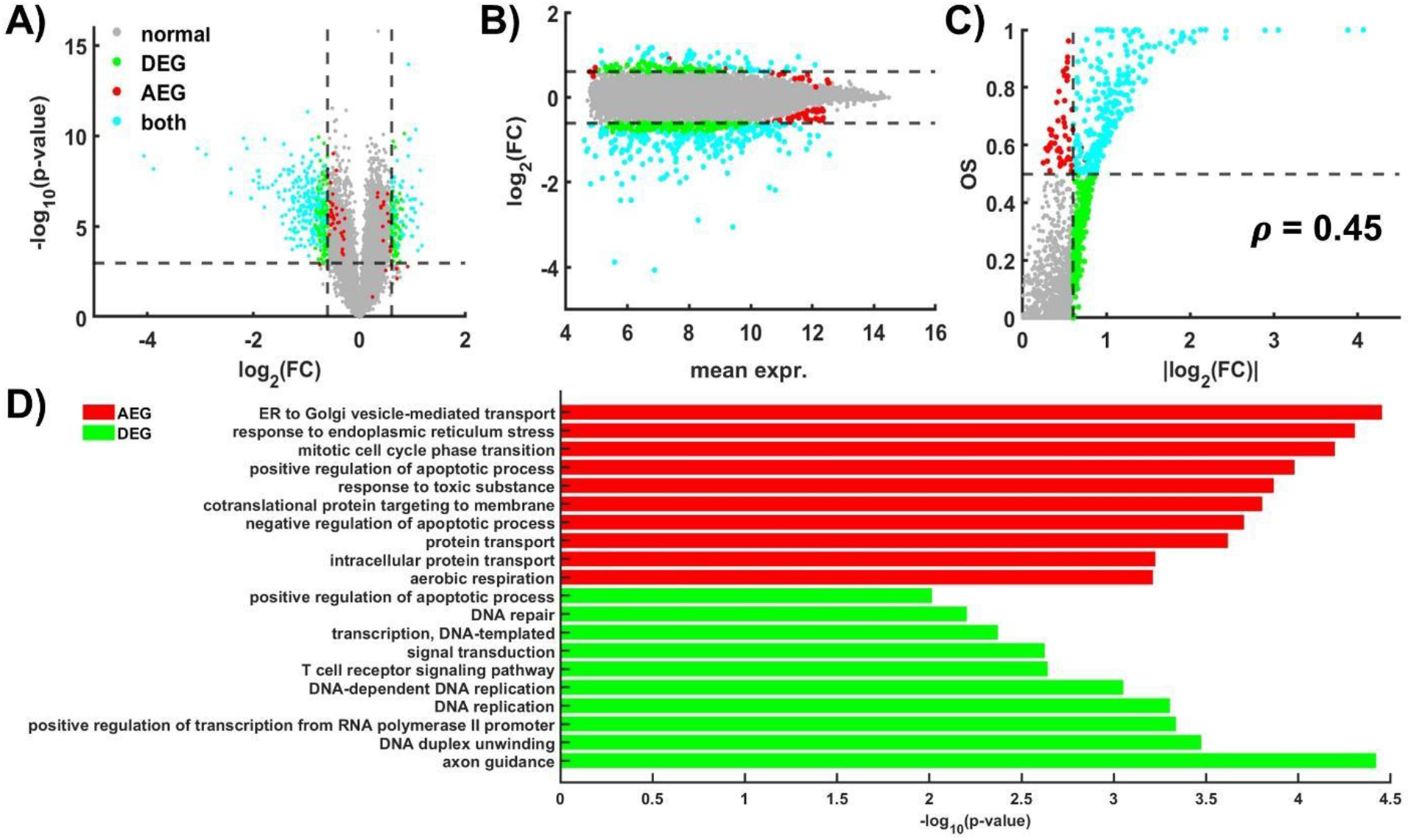
Comparison of MAGE and DEG on the breast cancer *γ* – T3 treatment profile. (A) Volcano plot based on the raw/uncorrected p-value of DEG and FC. (B) TPM vs FC scatter plot. (C) Relationship between FC and *OS*. Colors indicate if a gene is identified as DEG and/or AEG. (D) Top 10 GO terms enriched by AEGs and DEGs exclusively.

Next, we compared the biological significance of both DEGs and AEGs (Table S1) by GO enrichment analysis using DAVID on each gene set (Table S2 and Figure S13). The GO analysis showed a prevalence of terms related to cellular stress response, protein transport, gene expression regulation, and apoptotic processes in common. There is a large overlap between AEGs and DEGs, but some genes are identified exclusively in each set. To focus on the difference between AEGs and DEGs, we selected genes with low (<0.05) AE FDR and high (> 0.05) DE FDR as well as genes with low DE FDR and high AE FDR. Figure 5D shows the top 10 enriched terms based on the set of exclusive AEGs and DEGs, respectively. Notably, several of the top AEG-enriched terms are related to the ER stress response as a mechanism of tumor progression in breast cancer [33].

### Analysis of *mus musculus* mTor knockout RNA-seq data

To test the performance of MAGE with highly different gene expression profiles, we used mouse RNA-seq profiles (GSE134316) from healthy and mTOR knockout samples taken from mouse bone marrow [30]. The mechanistic target of rapamycin (mTOR) is one of the master regulators for growth and nutrient signaling, which regulates the global transcriptome [34]. Because of the categorization of mTOR as a master regulator, we expected to see a significant change in the expression of many genes between the two sample conditions. Figure 6 shows the expression profile with and without mTor, and as expected, there was a large variation in gene expression between the two conditions (Figure 6A) as well as an increase in the reported FDRs from MAGE (Figure S9B). Since there are a larger number of genes with high FC and moderate expression levels, the CER is stretched to encompass many of the genes that are often identified as DE (Figure 6B-C).

**Figure 6.**
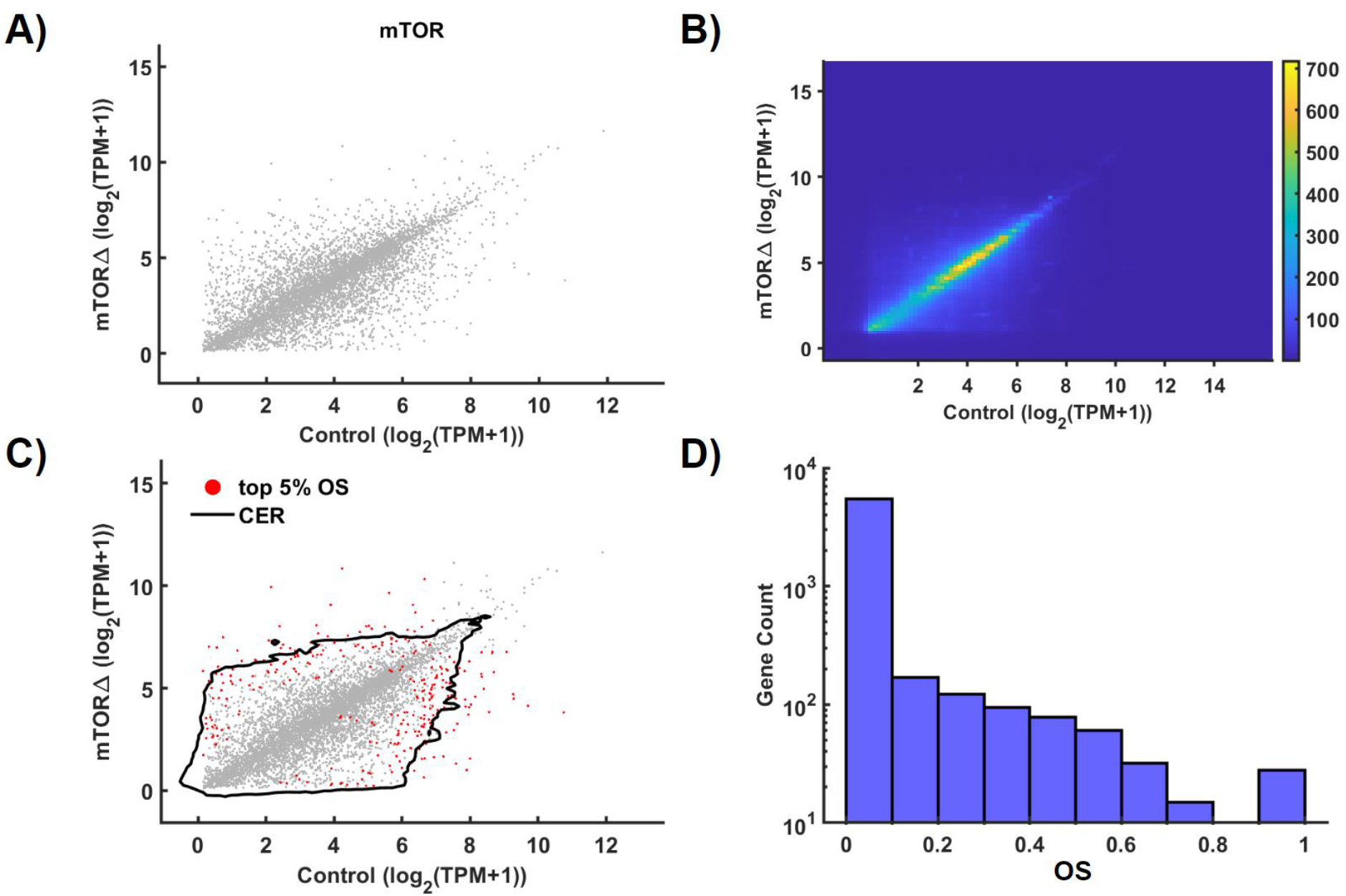
MAGE applied to the mTOR KO mouse profile. (A) Mean RNA-seq TPM levels of each gene per sample type after filtering. (B) CPDF from all genes. (C) CER based on the CPDF. (D) The distribution of *OS* from 1,000 MC sampling per gene.

Figure 7 compares AEGs and DEGs (Table S3), which shows significant agreement between the methods. There is still a positive ∼0.47 correlation between FC and *OS* (Figure 7C), which is similar to the ∼0.45 correlation in the *γ* – T3 profile. DEGs were also assessed using edgeR (Figure S14). To assess biological significance, we performed GO analysis on AEGs and DEGs (Table S4 and Figure S15). We found the set of AEGs to be enriched for genes related to cytoplasmic translation, cellular respiration, and oxidative phosphorylation (Table S4). DEGs were enriched in aerobic respiration and mitochondrial ATP synthesis (Figure 7D) as mTOR regulates both general and preferential mRNA translation [35] as well as the mitochondrial energetic adaptation [36]. These results suggest that MAGE can find genes from conventional DE analysis while finding additional gene sets that may be missed by conventional methods.

**Figure 7.**
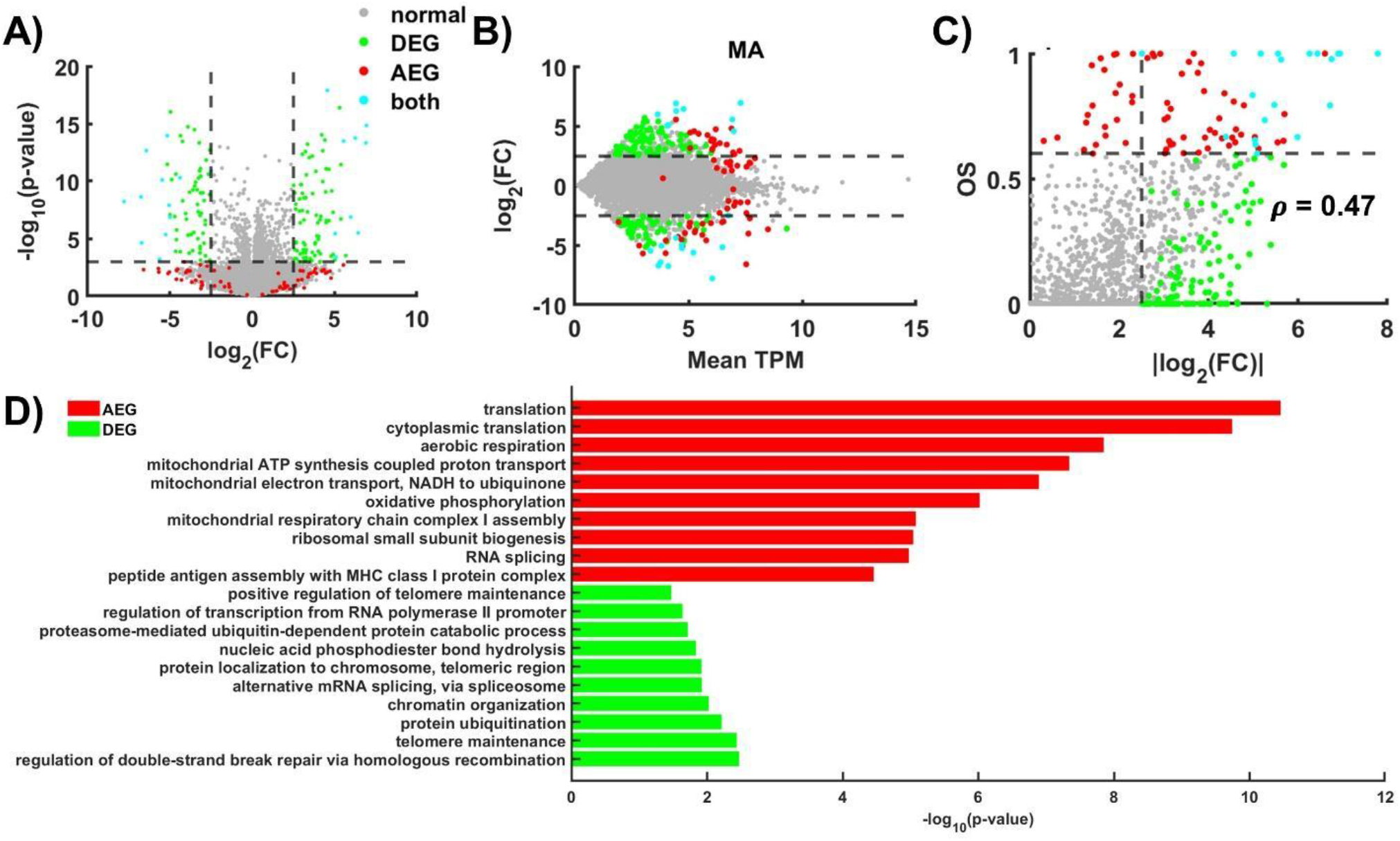
Comparison of MAGE and DEG on the mouse mTor KO profile. (A) Volcano plot based on the raw/uncorrected p-value of DEG and FC. (B) TPM vs FC scatter plot. (C) Relationship between FC and *OS*. Colors indicate if a gene is identified as DEG and/or AEG. (D) Top 10 GO terms enriched by AEGs and DEGs exclusively.

### Application of MAGE in scRNA-seq profiles

One of the emerging applications for RNA-seq is single-cell RNA-seq (scRNA-seq) which often exhibits high cell-to-cell variability. Profiles with large sample sizes are generally preferred, but an efficient method to identify DEGs or AEGs robustly with few samples is desirable. To compare DEG and AEG identification in single-cell data we analyzed the GSE98816 and GSE99235 profiles of mural and endothelial cells isolated from mouse brain and lung tissues [37,38]. MAGE parameters were set to default values with the target containment of 0.95, grid density of 100, and 5 contours per iteration. The expression’s upper and lower 2.5% extremes were disregarded for OS quantification. The mean expression within brain and lung cells is shown in Figure S16, along with the determined CER and genes within the highest 5% of OS values. We noticed that the variance within these profiles is significantly larger compared to the previously examined bulk samples, which was expected. The resulting CER is quite large and encompasses the vast majority of gene mean values, lowering the overall OS distribution (Figure S16). The correlation between OS and FC is still present but slightly reduced compared to the previously analyzed bulk samples (Figure S17 B). Interestingly, the genes with the highest OS tend to be found near the outer periphery of the volcano plot (Figure S17 A). We observe that higher expressed genes tend to have higher OS values (Figure S17 C), as seen in the previous microarray and RNA-Seq profiles.

Next, we asked how the sample sizes affect the robust AEG identification and compared its performance with that of conventional DE analysis. A greater number of samples is always better but with a diminishing return on preparation costs. Previous studies analyzing bulk profiles have noted that the optimal sample size is highly dependent on the variability of the data and in cases of low variability relatively few samples (<10) are necessary for reliable results [39]. Since scRNA-seq profiles tend to have significantly higher variability, we expected to see the need for a higher number of samples to provide consistent predictions. To test how many samples are required for robust AEG and DEG identification, we performed a random sample permutation in both of the scRNA-seq profiles and assessed the change in the OS distribution and the CER used to quantify OS. To quantify CER similarity, we used an MC area estimation to find the area overlap between the original CER (using all samples) and the reduced CER (using a subsampling) using equation 15 (Figure S18 A). Then we calculated the fraction of overlap between the top 5% AEGs and DEGs from subsamples and entire samples (*N*_*tot*_ ∼ 3000), which is a function of the number of subsamples. To consider the variability of these metrics we performed 4 trials of subsampling.

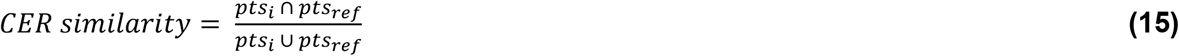

Figure S18 shows that the MAGE performance is saturated with about 100 samples in both the CER similarity and the mean OS stagnated before 100 samples. This demonstrates that for this particular data, MAGE requires around 100 samples to get the optimal CER. Comparing the 5% classification consistency graphs (Figure 8), we notice that genes classified as AEGs tend to be much more stable with 50 samples (73% +/- 6%) compared to both FC (52% +/- 9%) and p-value (40% +/- 5%) (DEGs). To examine this further we focused on the range of 20-500 samples (Figure 8) and noticed that OS classification reaches 75% +/- 5% consistency with only 40 samples (Figure S18 B), while genes classified by FC or p-value do not reach similar consistency until 200 samples are used. This suggests that MAGE results are robust with subsampling, and AEGs may be efficient with small numbers of scRNA-seq profiles.

**Figure 8.**
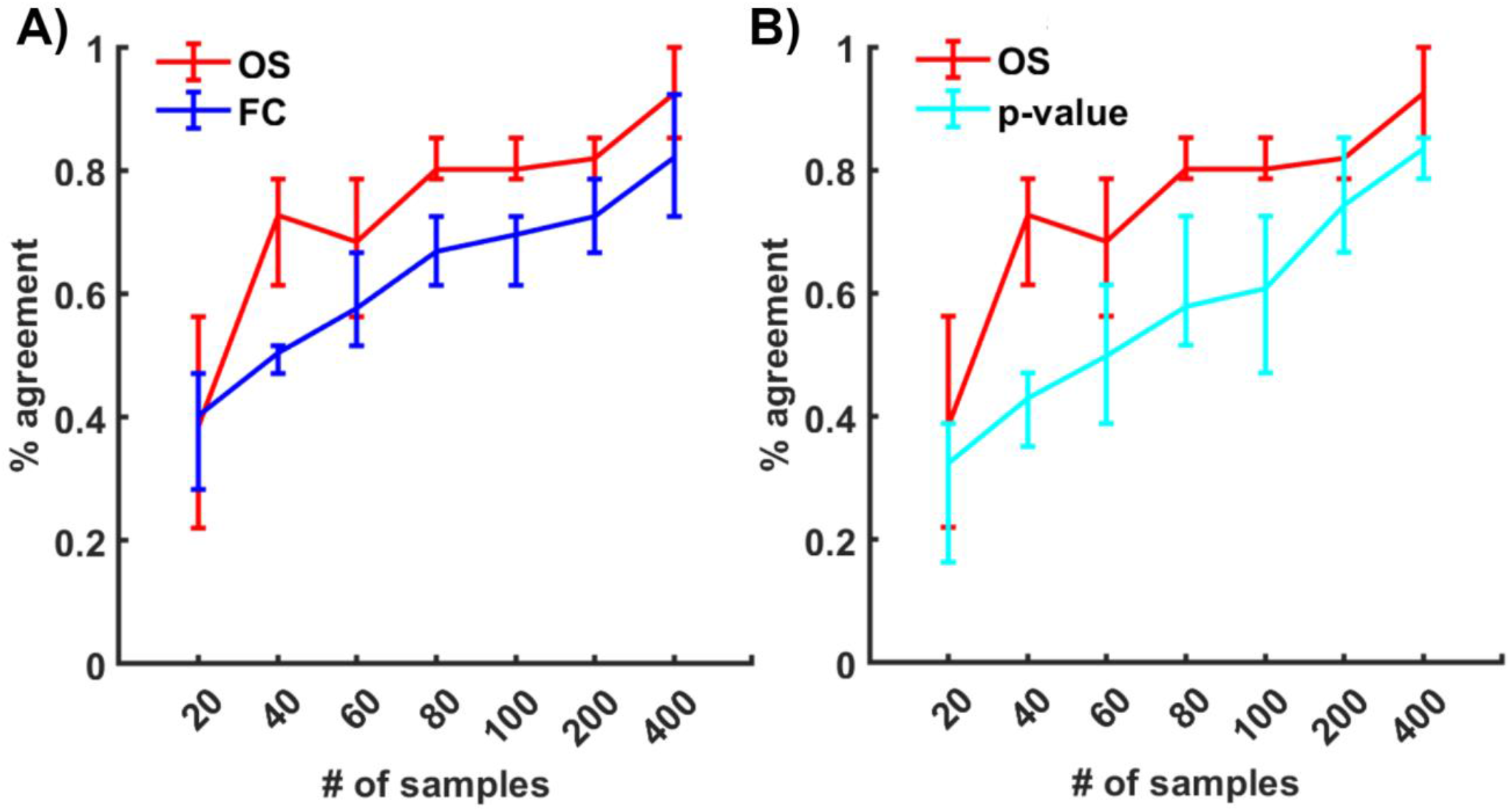
Robustness of AEG and DEG identification by the number of samples. (A) Robustness of AEG and DEG identification between brain and lung mural cells by the fraction of overlap of the top 5% genes at the indicated sample number and all samples for AEG (red) and DEG (blue), respectively. Lines represent mean values over the 4 trials, and error bars represent the maximum and minimum out of the 4 trials. (B) Consistency of genes determined by the lowest 5% p-value (cyan) and highest OS (red).

To test whether identified AEGs are biologically relevant, we performed the GO enrichment of AEGs and compared it to that of DEGs. For both the mural cells and endothelial cells, we identified exclusive AEGS (exAEGs) by sorting for genes with an OS greater than 0.1 and an FC less than 2. Here the exAEGs represent the set of genes identified by AE but missed in conventional DE approaches. We also sorted for exclusive DEGs (exDEGs) by selecting genes with an OS below 0.1 and an FC above 2.5. These thresholds selected roughly 200 genes for each list. Table S5 shows the pathway enrichment results for exAEGs, while the exDEGs did not provide significant (FDR < 0.05) enrichment results. This supports that AE can identify functionally relevant genes that are missed by DE analysis, while also retaining the most significant functional DEGs. We investigated prominent exAEGs (high OS and low FC) and one of the clear examples is CXCL12 (OS = 0.36 and FC = 0.70), which plays a significant role in brain development, particularly through angiogenesis [40].

## Discussion

### Interpretation of MAGE expression

It is important to note that differentially expressed genes and aberrantly expressed genes are different. Differential expression implies significant up/down-regulation of a gene which is usually measured by the logarithmic fold-change. Here we define an aberrant expression as a deviation from the group. Therefore, our main assumption is that the majority of genes should not be considered gene signatures and that the genes of biological interest will be found further from the majority. The sets of DEGs and AEGs in many cases will contain significant overlap. However, depending on the types of samples being studied these sets may be significantly different. Both differential expression and aberrant expression may provide potential insight into the features responsible for the biological variation of samples. Therefore, MAGE is not intended to be a replacement or improvement to differential expression analysis, but to be used as an alternative analysis, particularly in cases where differential expression may not be the most informative.

We have also assumed a log-normal distribution for the mean expression of a gene across samples. Although this may not capture the long-tailed nature of some genes, we use this assumption to construct a probabilistic landscape of the cumulative probability distributions from every gene in the experiment.

### Potential application in single-cell sequencing

Alterations in the transcriptional programs that take part in the onset and progression of diseases can go unnoticed when bulk tissue samples contain highly heterogeneous mixtures of cell types. Recent studies have utilized flow cytometry and subsequent RNA-seq to examine transcriptional evolution throughout disease progression and identify biomarkers that only occur in individual cell types [41]. However, scRNA-seq introduces many difficulties in generating informative and consistent results. This is because read counts taken from single-cell profiles are notoriously low which can often produce a high expression bias in the DE classification [42]. Furthermore, in the commonly used parametric methods for DE analysis, the assumptions necessary for reliable results are often not met in single-cell studies. Stochastic switching between gene network ‘on’ and ‘off’ states becomes more apparent at the low levels of RNA species present in most single-cell profiles. This leads to bi-modal behavior exhibited in scRNA-seq that is often unobserved in bulk [43]. MAGE is designed to work well in the presence of bi-modal expression distributions and can easily identify genes found between states. The ability of MAGE to identify aberrant genes without the need to verify prior distribution assumptions would seem to make this a promising method for scRNA-seq analysis.

## Conclusion

We have presented a novel methodology aimed at addressing the limitations of conventional DEG analysis. By analyzing the expressional probability overlap between the genes of 2 samples, our method offers a unique perspective, focusing on the identification of genes exhibiting aberrant expression patterns rather than solely examining differential expression. Through extensive validation using diverse datasets, we demonstrated the robustness and applicability of this approach across various experimental conditions related to cancer.

This methodological shift towards evaluating gene expression based on the deviations of genes relative to the entire profile offers promising insights into understanding disease mechanisms. The ability to identify functionally relevant gene signatures, those exhibiting aberrant expression patterns, presents opportunities for novel biomarker discovery.

Furthermore, our approach’s adaptability for large-scale omics datasets while remaining computationally feasible enhances its potential for widespread application in diverse biological studies. However, while our method showcases significant promise, there remain avenues for further refinement and validation. Future research should focus on refining the algorithm, particularly its adaptability to single-cell sequencing data, and exploring its utility across various disease contexts beyond cancer. Additionally, continued efforts in validating identified gene signatures and their biological relevance will be essential for translating these findings into clinical applications.

In essence, MAGE represents a complementary framework for identifying aberrant gene expression, which can run in parallel to the identification of DEGs. Often MAGE can return consistent results in ambiguous scenarios where assumptions are uncertain, and its potential to unveil biologically significant gene signatures holds promise for advancing our understanding of disease mechanisms and fostering the development of precision medicine strategies.

## Acknowledgments

RJ was supported by Institutional Research Grant IRG-18-159-43 from the American Cancer Society. We are grateful to all the Joh lab members for the critical reading and helpful discussions.

## Author Contributions

MB and RJ perceived experiments. MB designed models and performed the analysis. MB and RJ wrote the paper.

