## Supplementary Information for "MAGE: Monte Carlo method for Aberrant Gene Expression"

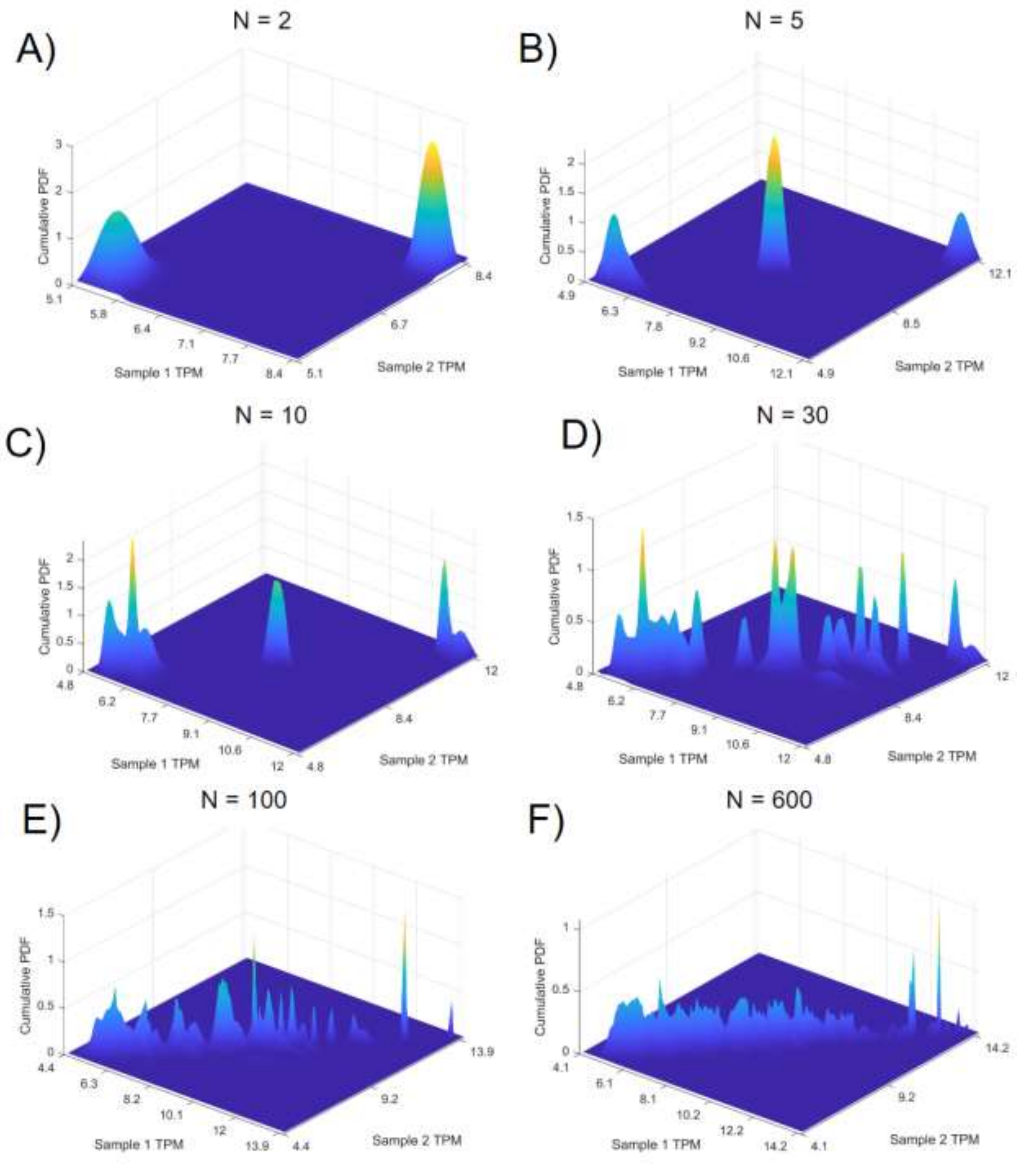


**Figure S1. Cumulative PDF as a function of the number of genes.** Surface plots of the probability density matrices formed by running the topological analysis of the breast cancer $\gamma$-T3 treatment profile (GSE21946) with indicated number of selected genes.

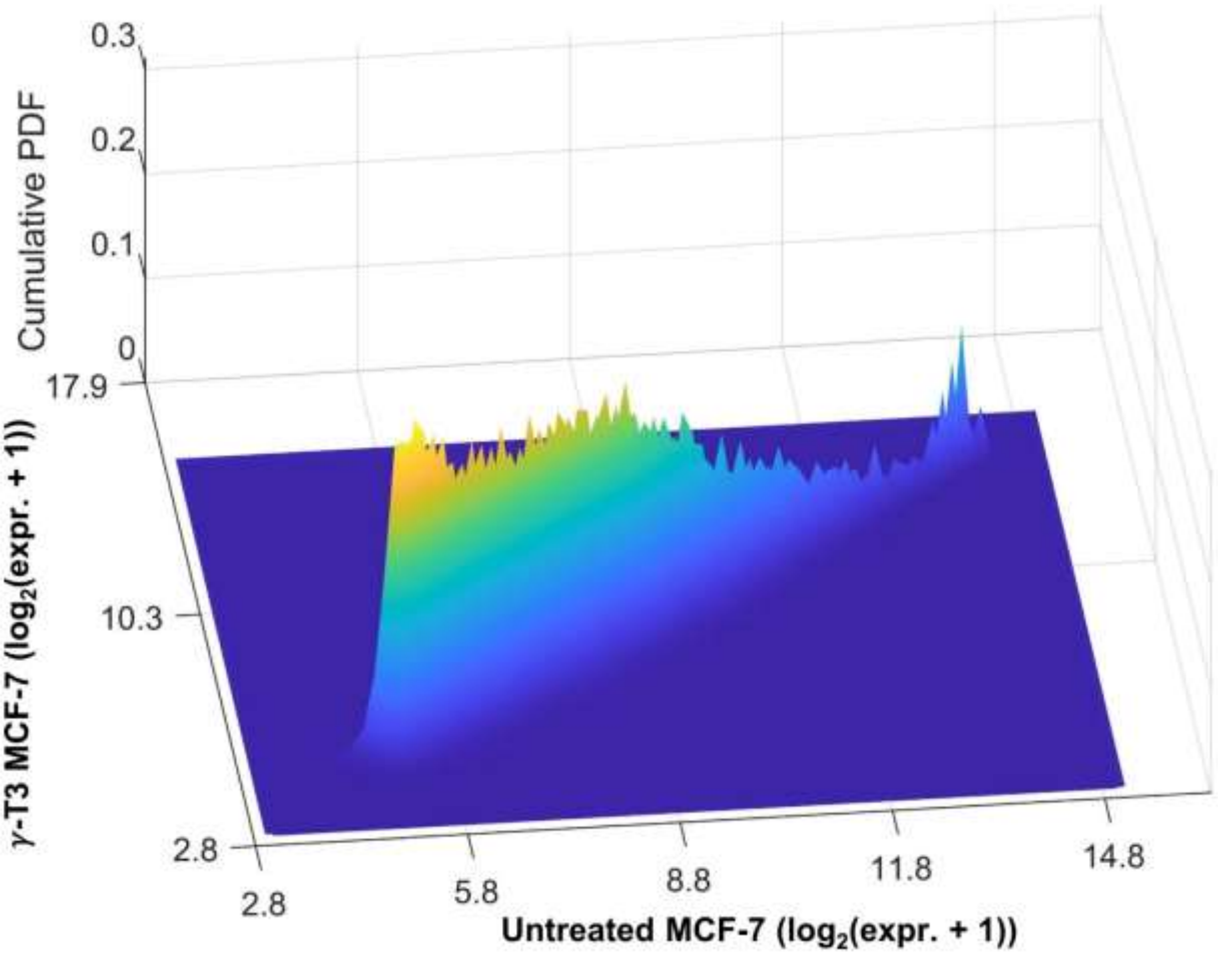


**Figure S2. Cumulative PDF from the breast cancer $\gamma$-T3 treatment profile against control (data from GSE21946).**

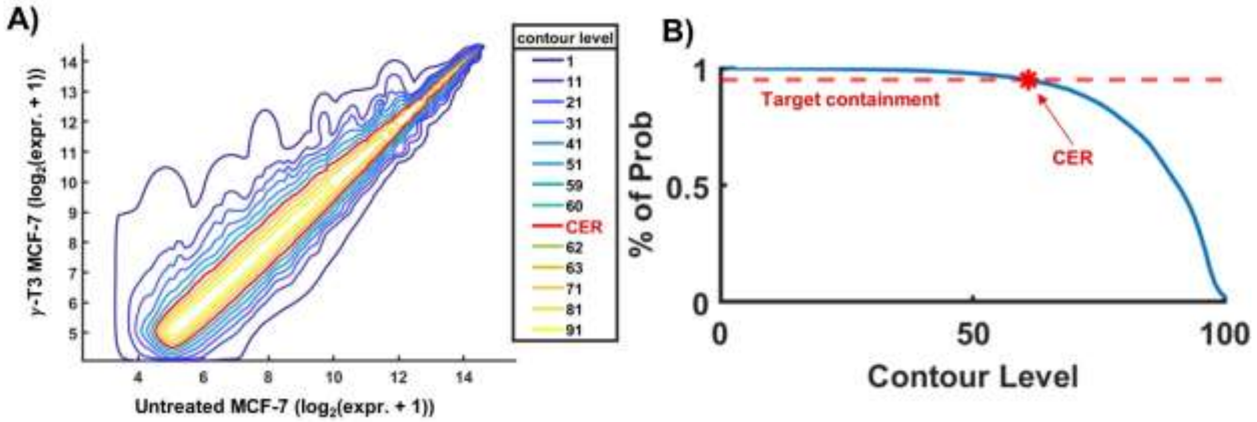


**Figure S3. Probability containment of CPDF contours and selection of CER.** (A) Representative contours as indicated by color were selected at various heights/levels of CPDF using the breast cancer $\gamma$-T3 treatment profile. The optimal contour selected as the CER is shown in red. (B) The fraction of contained genes at each contour level.

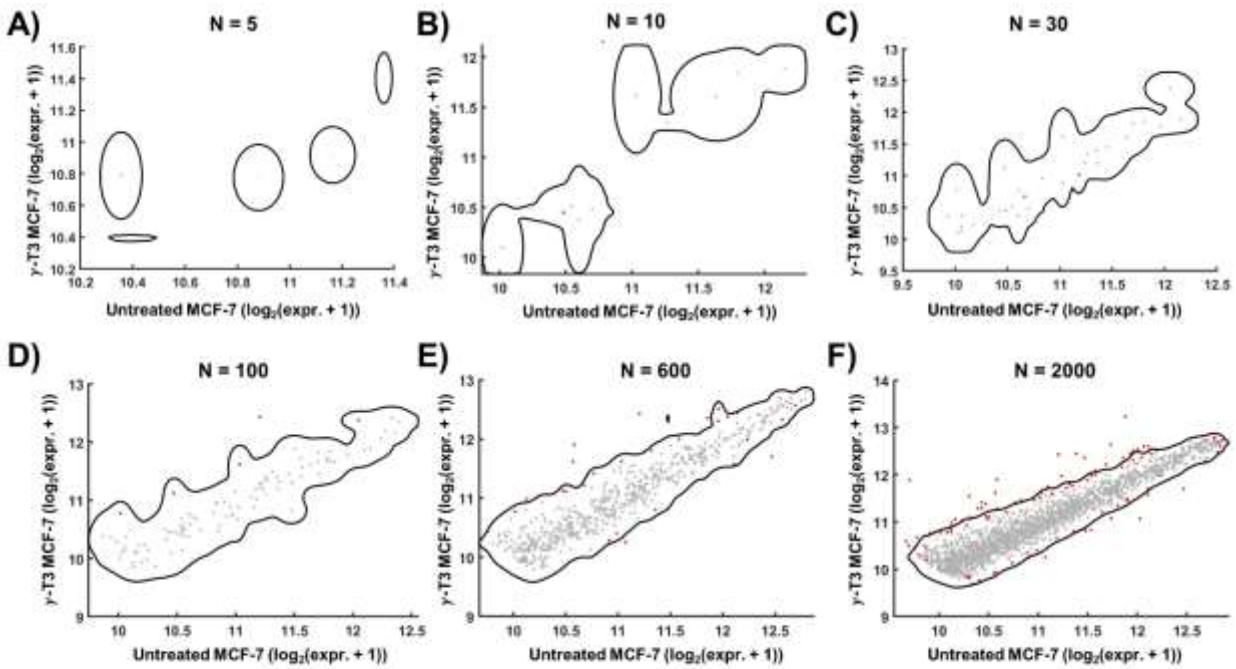


**Figure S4. CER with different number of genes using breast cancer $\gamma$-T3 treatment profile.** The indicated number of genes was randomly selected. Data points represent mean expression values for individual genes and genes with the highest 5% of outlier scores displayed in red.  The black curve represents the CER boundary.

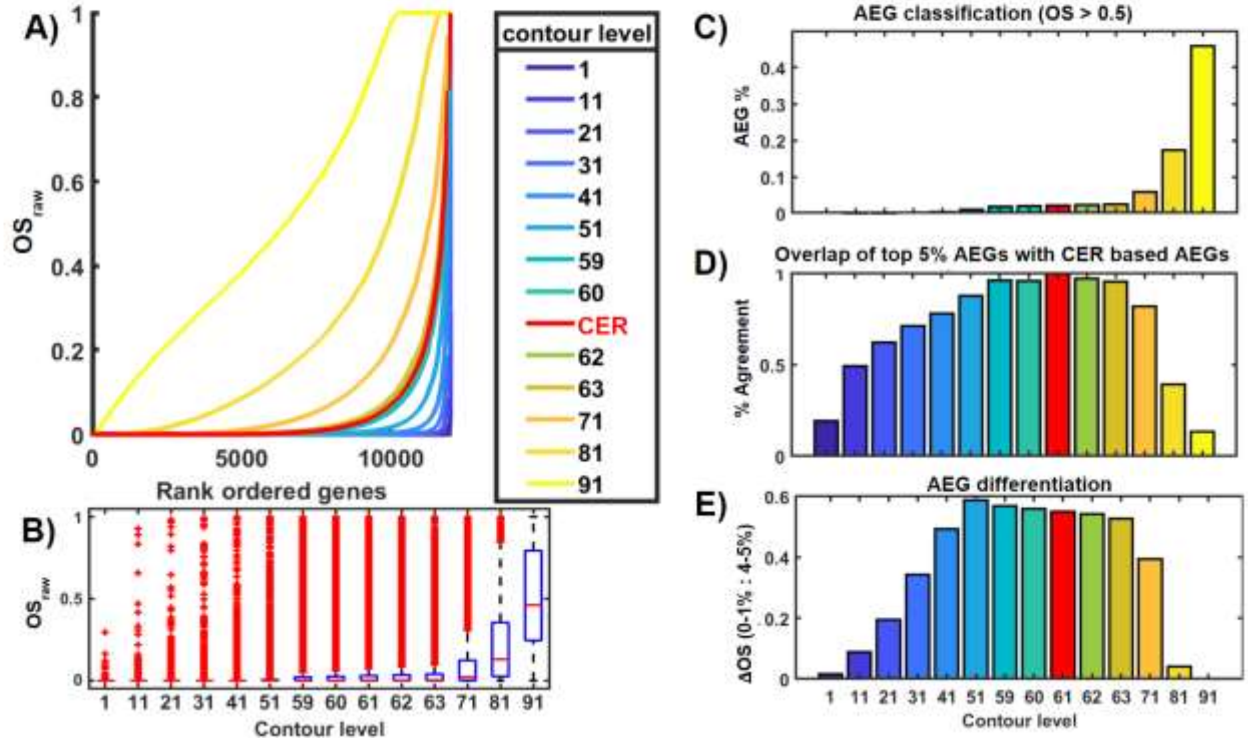


**Figure S5. Comparison of AEGs from CER and other CPDF contours.** (A) Distributions of $OS$ by indicated contour as the CER. Each curve represents the cumulative distribution of $OS$. (B) Distribution of $OS_{raw}$ based on each contour level. (C) AEG identification by MAGE using $OS$ as a cutoff ($OS$ < 0.5). (D) Overlap of top 5% $OS_{raw}$ genes between the optimal contour and indicated contours. (E) The difference of $OS_{raw}$ of genes in the highest 1% and 5% as a selectivity for AEGs (data from GSE21946).

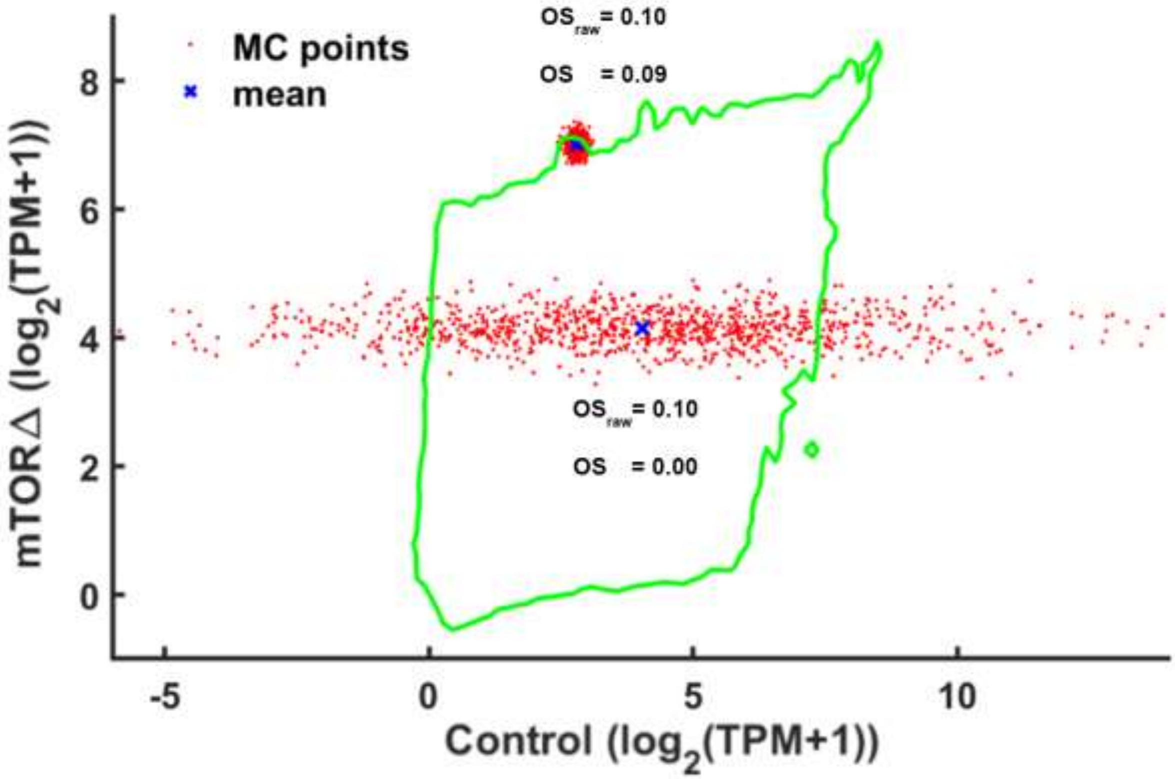


**Figure S6. The effect of mean expression and variance on $OS$.** Scatter plot of MC sampled points for two genes with similar $OS_{raw}{\sim}0.1$. The mean expression of each gene is shown in blue and 1,000 randomly selected points from each gene's PDF are shown in red. The estimation of $OS$ then adjusted by the distance from CER. Data are from GSE134316.

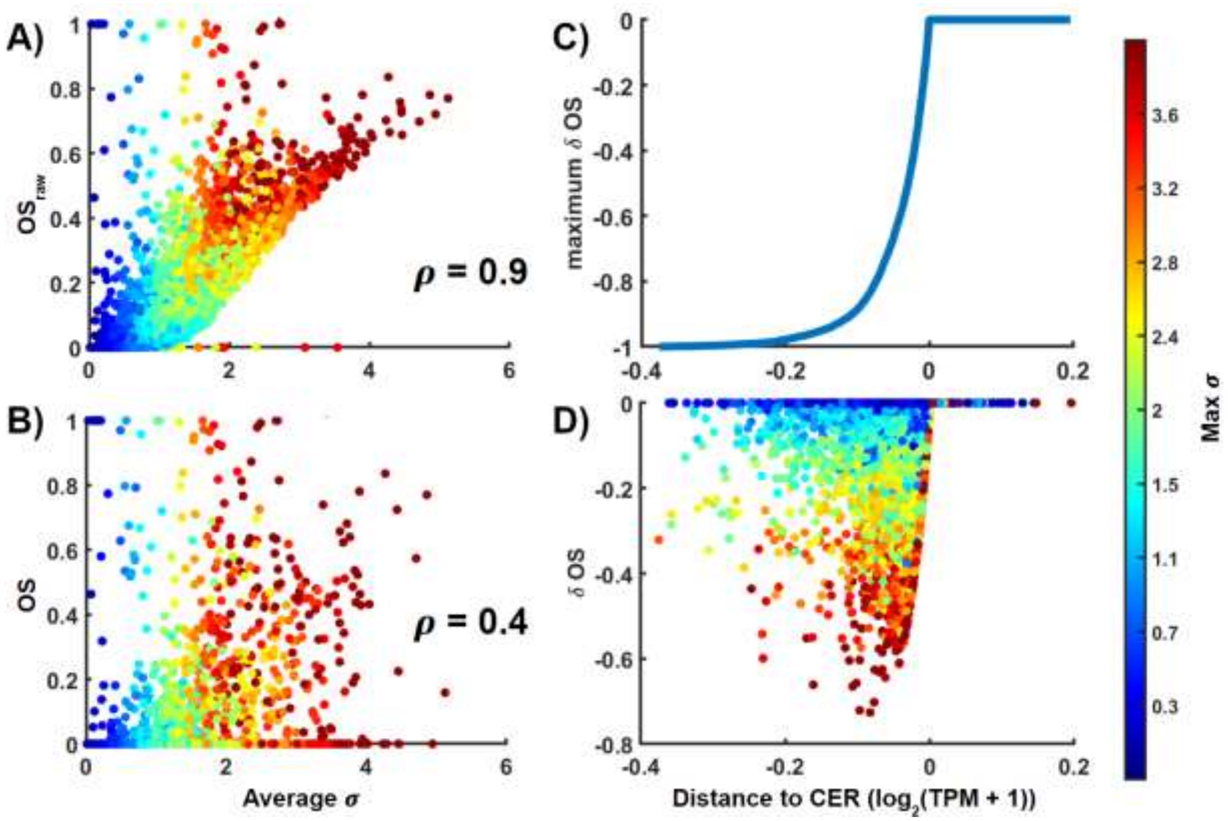


**Figure S7. Correction of $OS_{raw}$ by the distance to CER.** (A) Scatter plot of $OS_{raw}$ and SD (denoted as $\sigma = \sqrt{\sigma_x^2 + \sigma_y^2}$) of all genes in the mTOR KO profile. (B) Scatter plot of $OS$ and SD. ρ indicates the Pearson correlation. (C) The correction term $\delta OS = OS_{raw} - OS$ based on eq.10 as a function of the distance to CER. (D) Scatter plot of $\delta OS$ and mean SD. Color corresponds to the maximum SD of each gene calculated in both conditions.

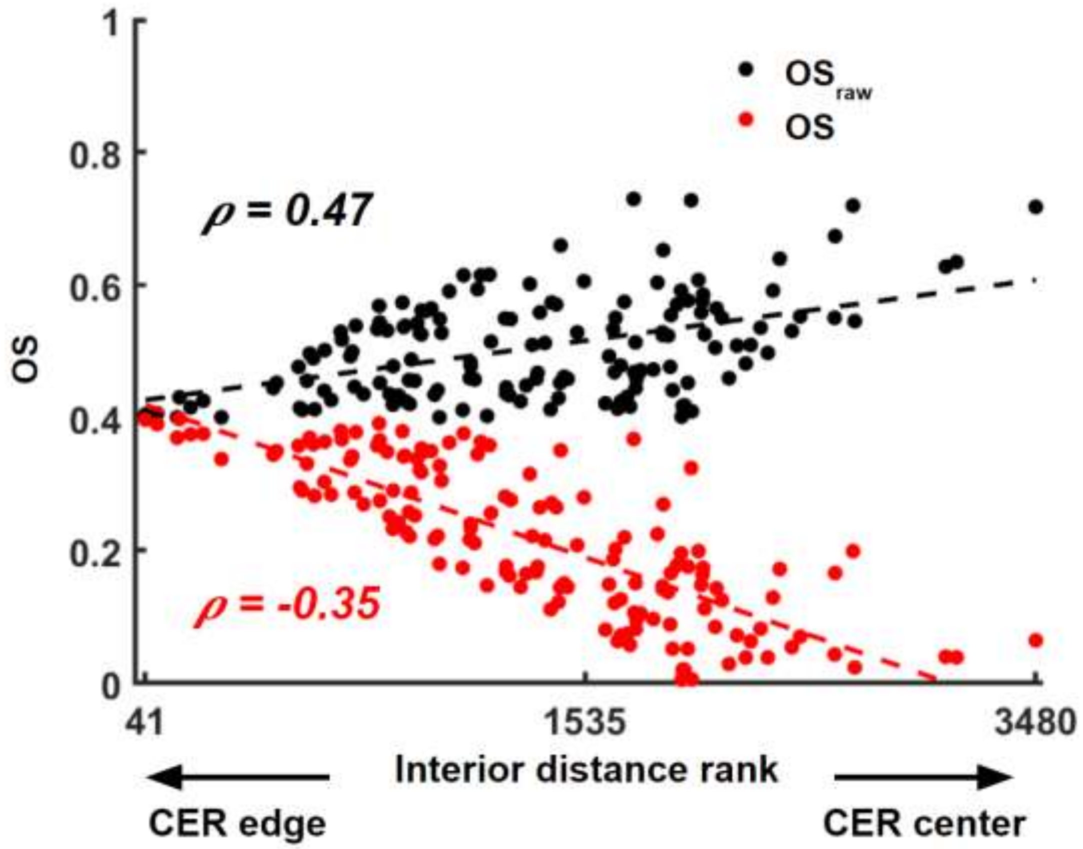


**Figure S8. $OS_{raw}$ and $OS$ as a function of interior distance.** $OS$ of Genes with **.** $OS_{raw} > 0.4$ and $OS > 0$ were plotted with the distance to CER (ranking by distance to CER). A smaller distance indicates genes are near the edge of the CER, while genes with large distances are found near the center.

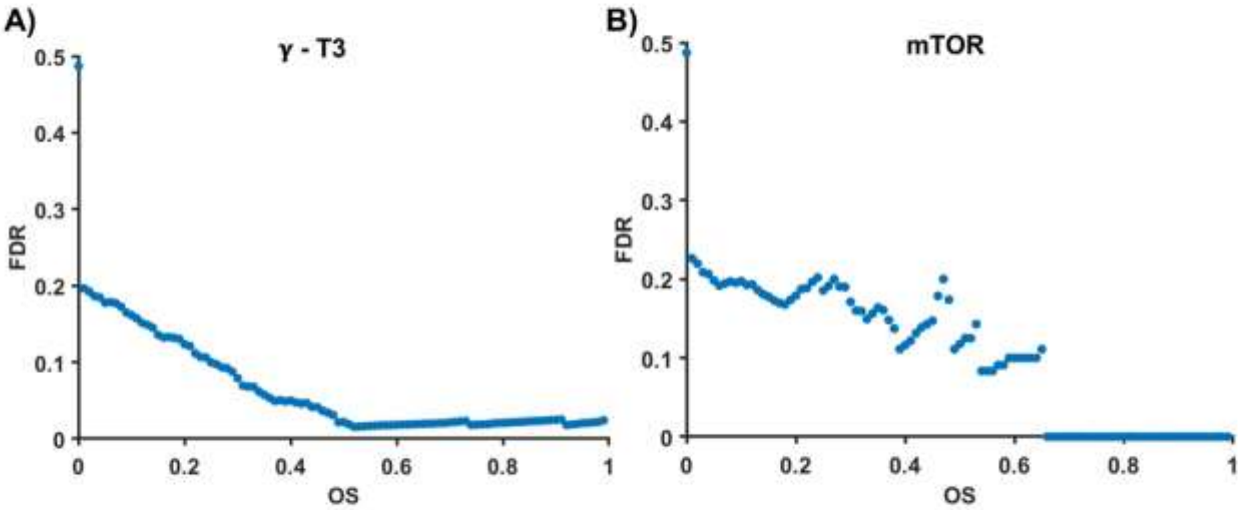


**Figure S9. FDR determined by sample permutation.** FDR was estimated (using eqs. 11 and 12) in both (A) breast cancer $\gamma$-T3 treatment and (B) mTOR KO mouse profiles.

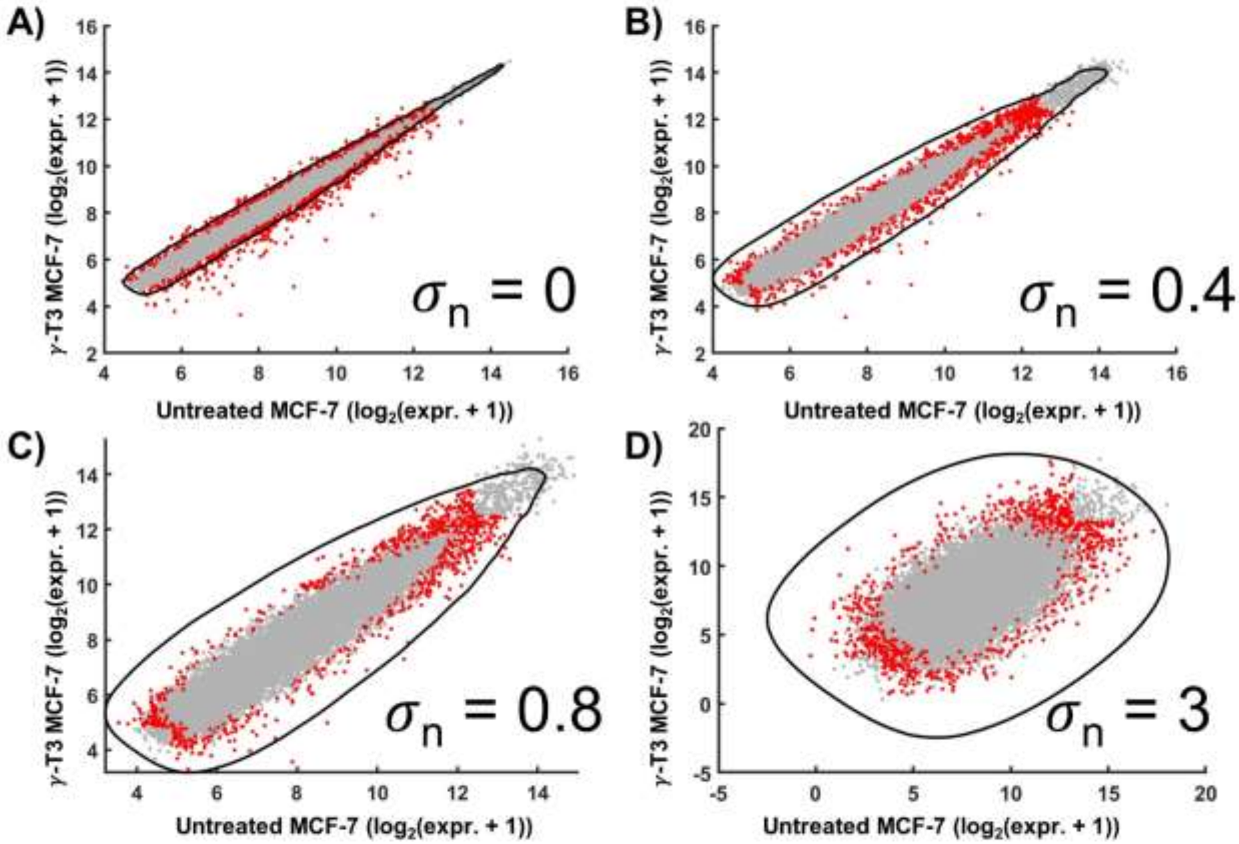


**Figure S10. Effect of Gaussian random noise on CER.** For each gene, Gaussian random noise with indicated SD was added to both x- and y-data. After adding normally distributed noise with varying SD $\sigma_n$. The mean expression of each gene is shown plotted in both conditions. Genes with the highest 5% of $OS$ values are shown in red. The CER from each profile is shown in green (data from GSE21946).

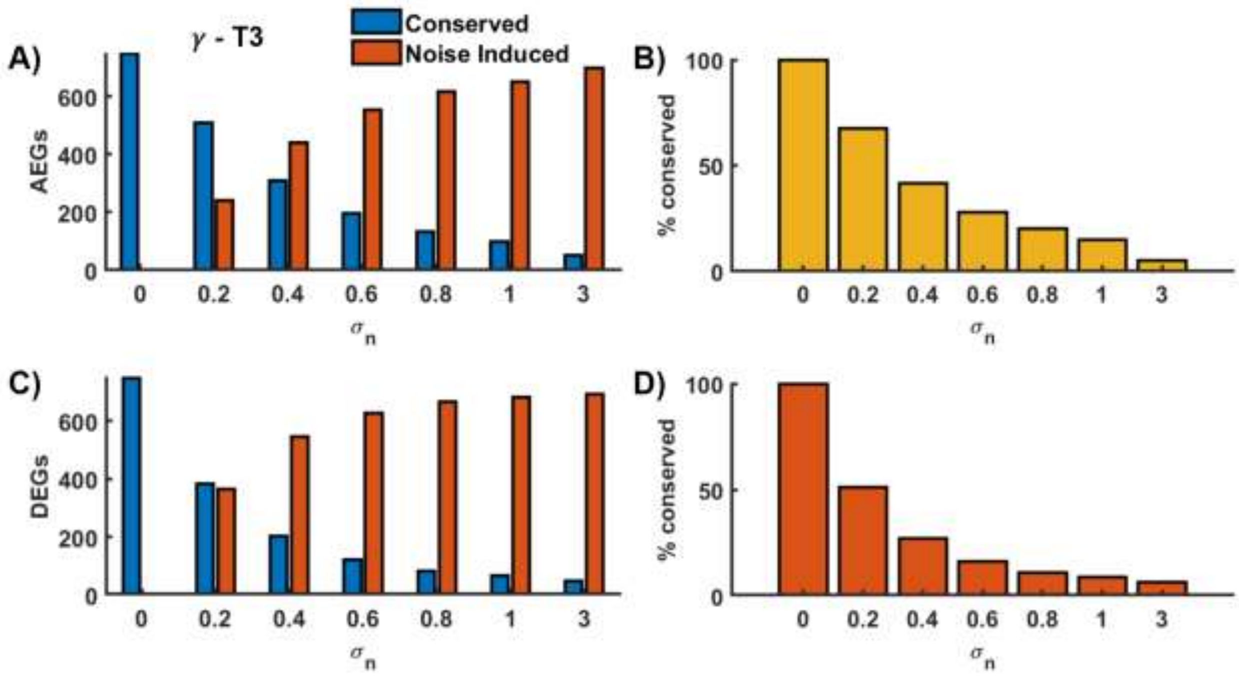


**Figure S11. Performance MAGE with varying levels of noise introduced in breast cancer $\gamma$-T3 treatment profile.** (A) The number of AEGs with indicated Gaussian noise. (B) Fraction of overlaps between no-noise-AEGs and AEGs with indicated noise. (C) The number of DEGs with indicated Gaussian noise. (D) Fraction of overlaps between no-noise-DEGs and DEGs with indicated noise (data from GSE21946).

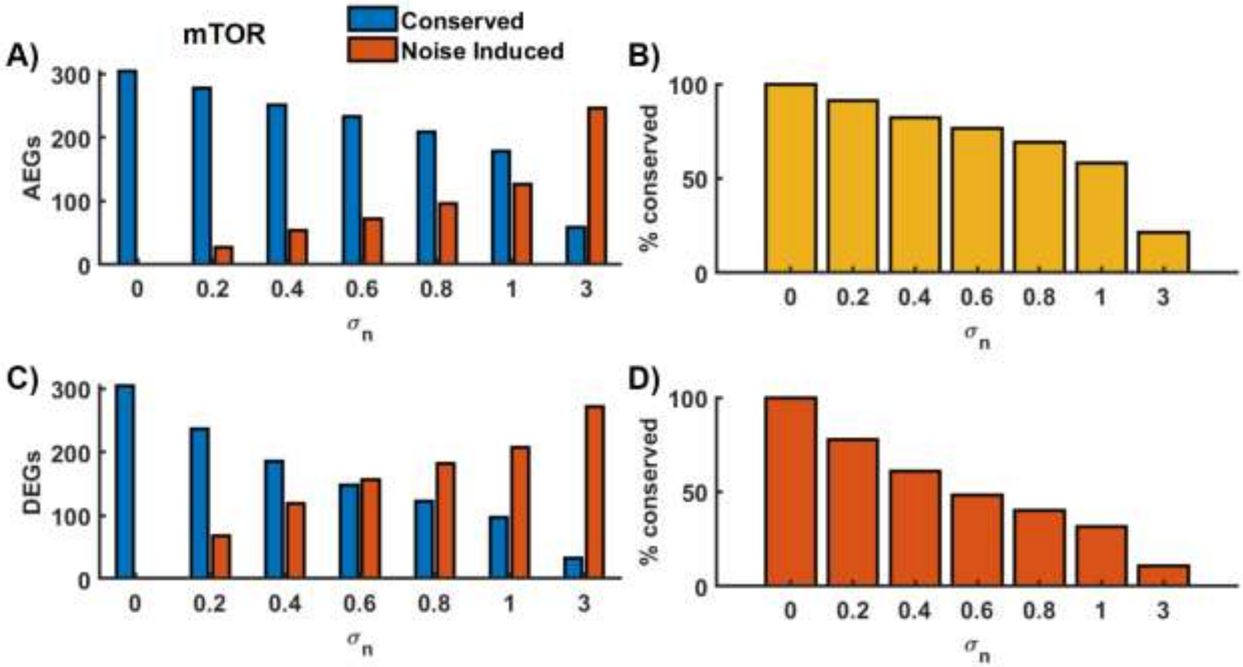


**Figure S12. Performance MAGE with varying levels of noise introduced in mTOR KO mouse profile.** (A) The number of AEGs with indicated Gaussian noise. (B) Fraction of overlaps between no-noise-AEGs and AEGs with indicated noise. (C) The number of DEGs with indicated Gaussian noise. (D) Fraction of overlaps between no-noise-DEGs and DEGs with indicated noise (data from GSE134316).

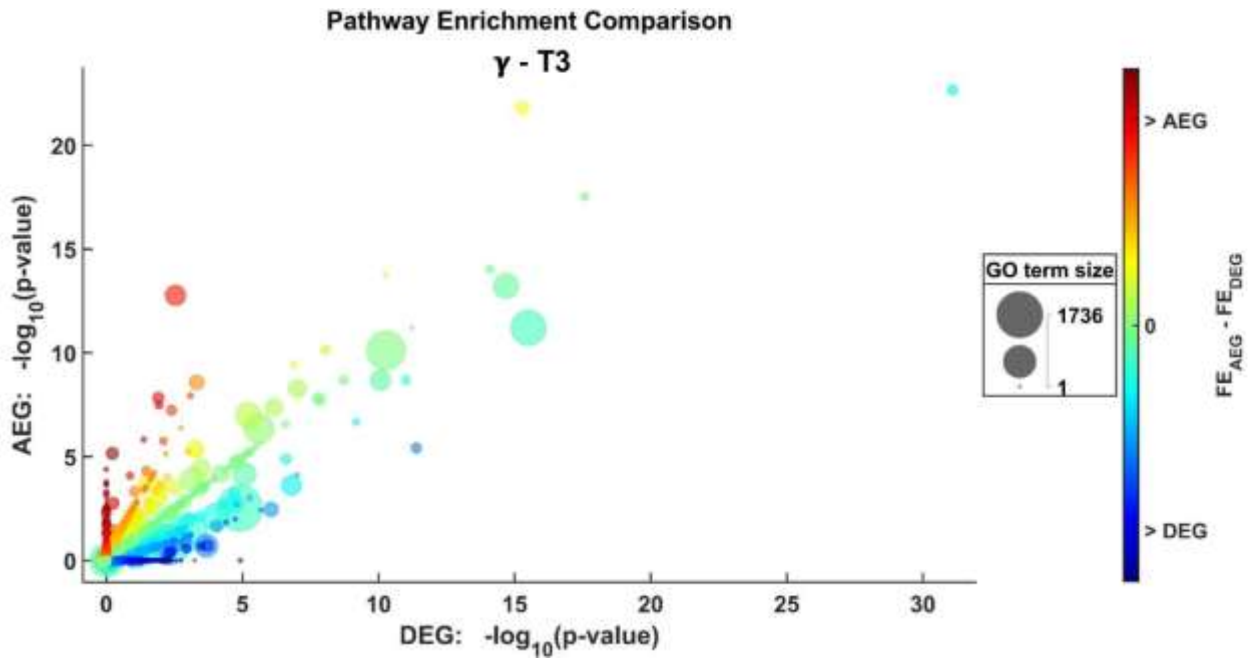


**Figure S13. Comparison of breast cancer $\gamma$-T3 treatment profile pathway enrichment of AEGs found using MAGE and DEGs found using t-test.** Bubbles represent individual GO terms. Size of bubble represents the total number of genes associated with the individual GO term from the DAVID *Homo sapiens* reference gene set. Color represents the difference in the fold-enrichment from the set of AEGs and DEGs.

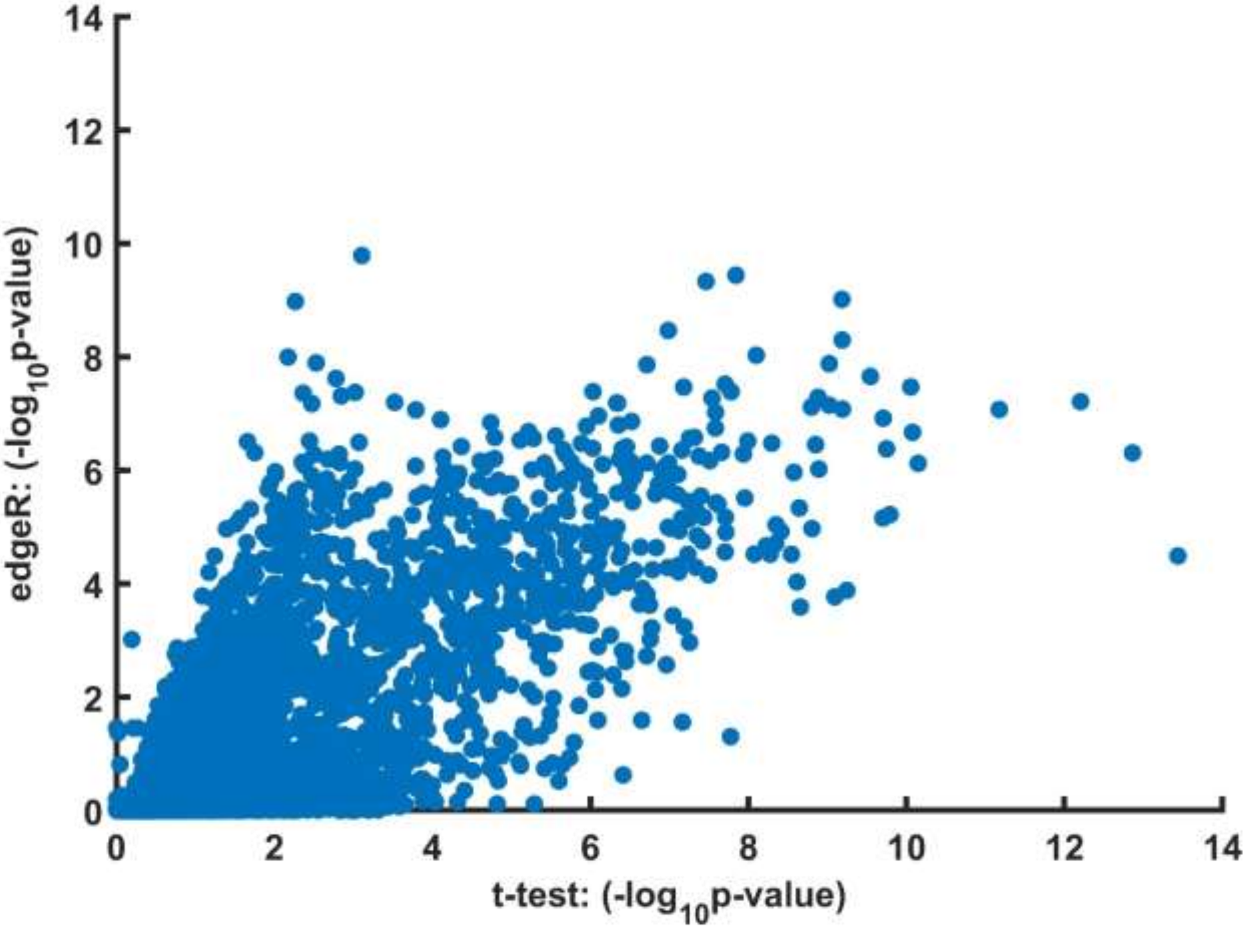


**Figure S14. P-value comparison using edgeR.** P-values determined by standard 2 sample t-test along the X-axis and by edgeR along the Y-axis (data from GSE134316).

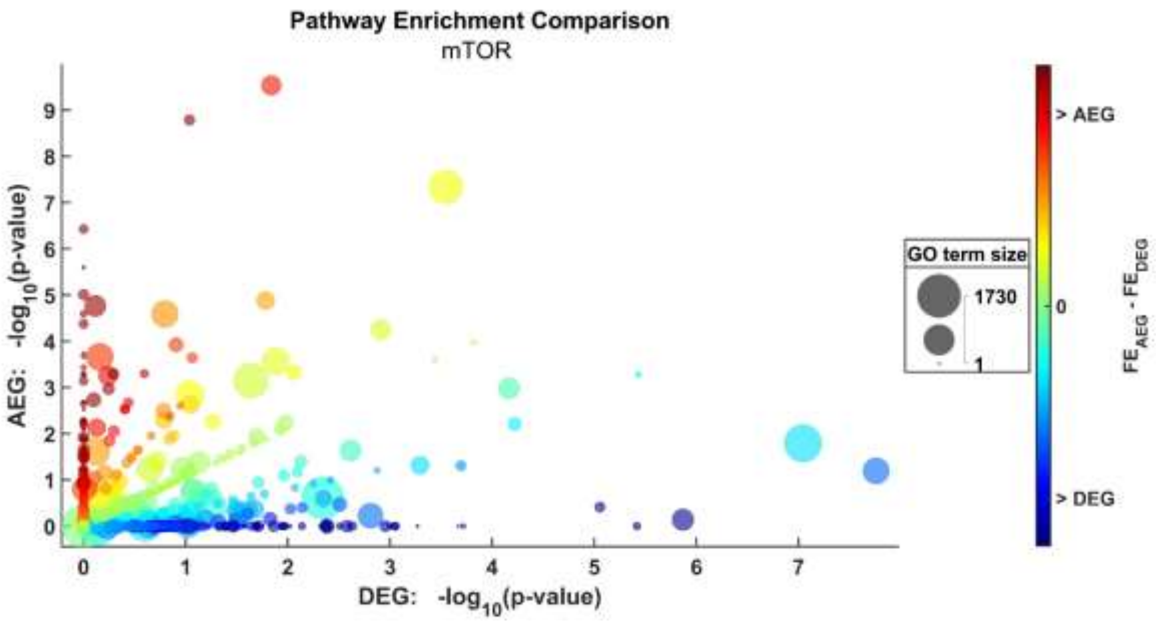


**Figure S15. Comparison of mTOR KO mouse profile pathway enrichment of AEGs found using MAGE and DEGs found using t-test.** Bubbles represent individual GO terms. Size of bubble represents the total number of genes associated with the individual GO term from the DAVID *Mus musculus* reference gene set. Color represents the difference in the fold-enrichment from the set of AEGs and DEGs.

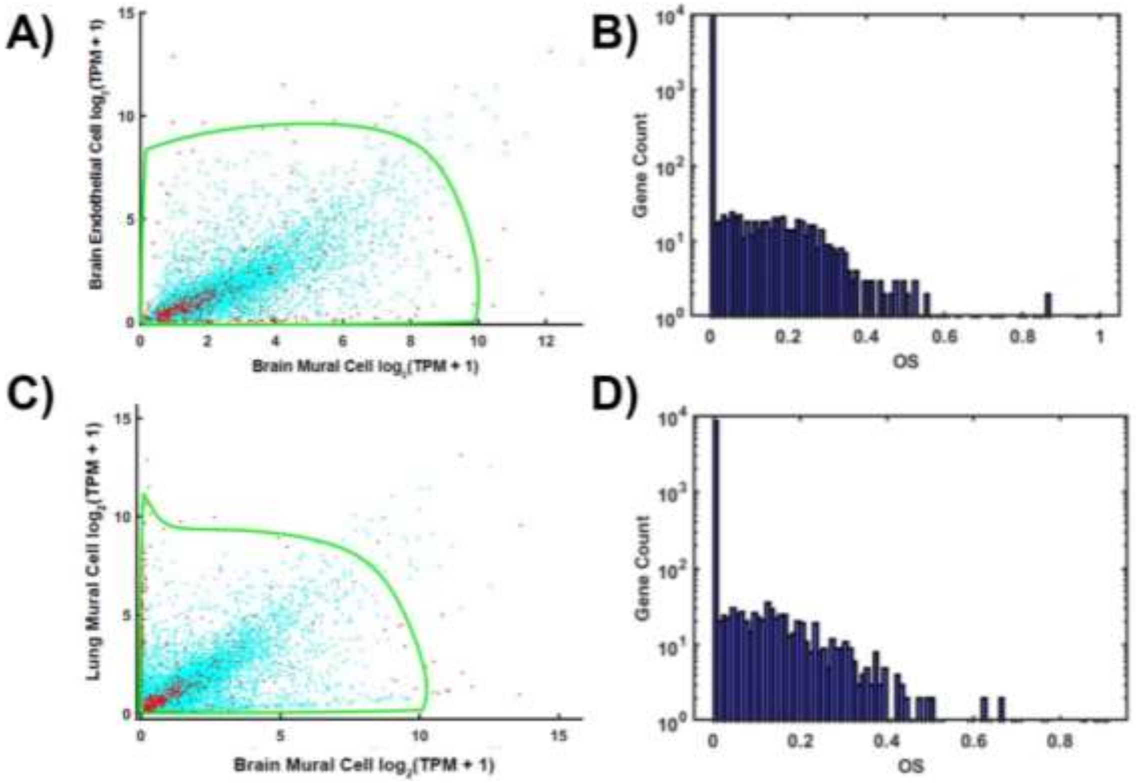


**Figure S16. Comparing brain mural, brain endothelial, and lung mural cells using scRNA-seq.** (A-B) MAGE analysis of brain mural and endothelial cells. (A) Mean expression of each gene and CER used to assess AE. (B) Distribution of OS. (C-D) MAGE analysis of brain mural and lung mural cells. (C) Mean expression of each gene and CER used to assess AE. (D) Distribution of OS.

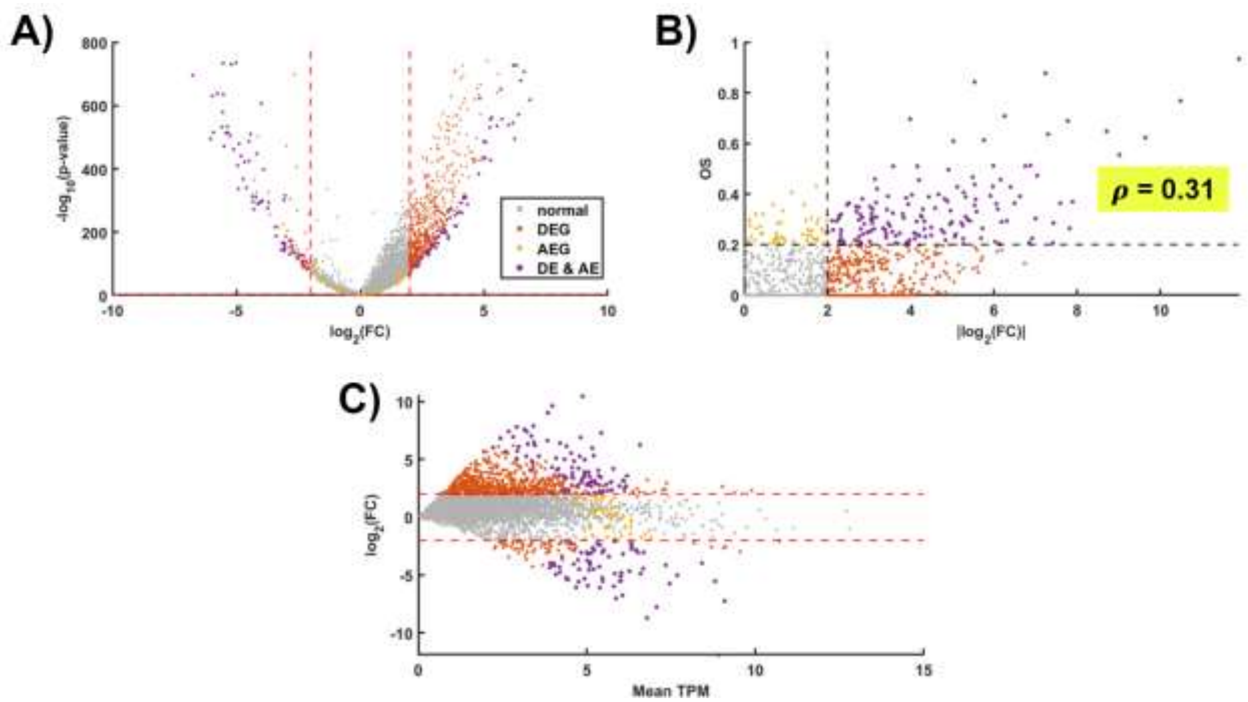


**Figure S17. Comparison of brain and lung mural cells.** (A) Volcano plot from the DEG analysis. (B) FC versus OS of all genes. (C) TPM versus FC of all genes. Color indicates normal genes (grey), AEGs (yellow) DEGs (red), and both (purple).

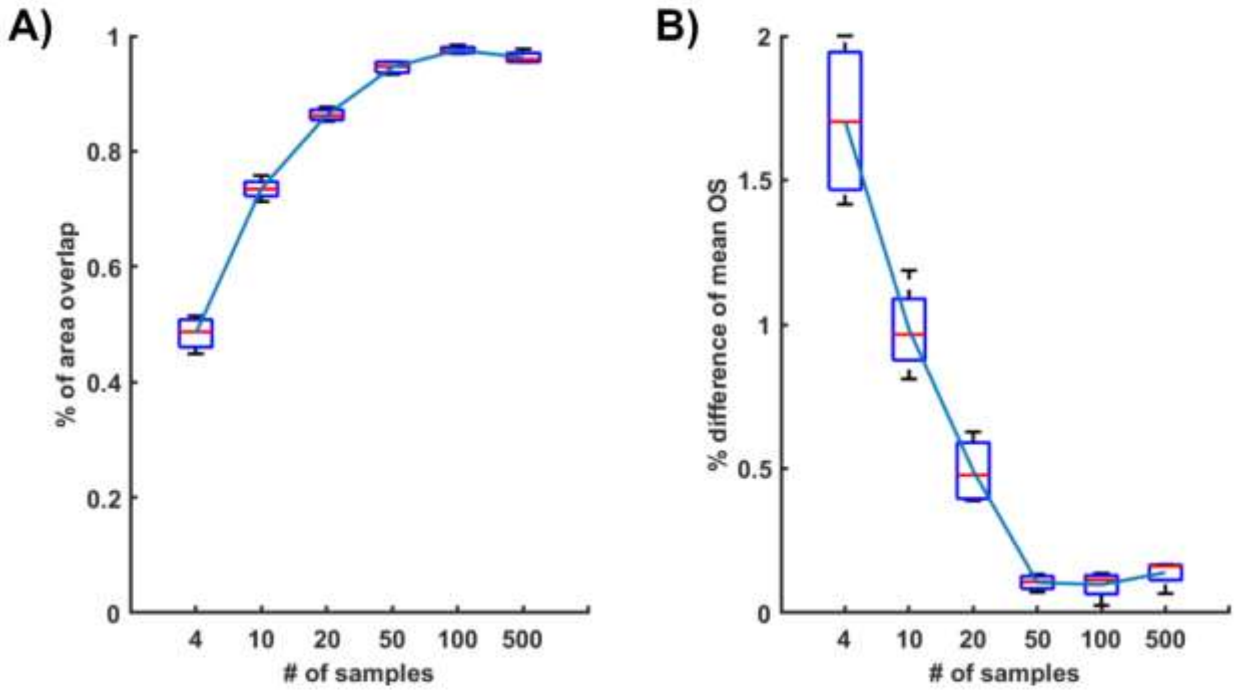


**Figure S18. MAGE performance consistency by subsampling.** (A) CER similarity in terms of the area of overlap between the reference CER (all samples) and the CER determined with the indicated number of randomly subsampled genes. The box plot was from 4 subsamplings. (B) Percent difference comparing the mean OS from the reference and with the indicated number of randomly subsampled genes. Data is brain and lung mural cells scRNA-Seq.
